# Social Exclusion Amplifies Behavioral Responses to Physical Pain via Insular Neuromodulation

**DOI:** 10.1101/2025.05.09.653162

**Authors:** Caroline Jia, Andrea Tran, Faith Aloboudi, Ella Say, Nick Thao, Christopher R. Lee, Kanha Batra, Amanda Nguyen, Aneesh Bal, Nathaniel N. Nono, Jeremy Delahanty, May G. Chan, Laurel R. Keyes, Reesha R. Patel, Romy Wichmann, Felix Taschbach, Yulong Li, Marcus Benna, Talmo D. Pereira, Hao Li, Kay Tye

**Author notes:** To Whom Correspondence Should be Addressed: Kay M. Tye, Ph.D. HHMI Investigator, Wylie Vale Professor, Salk Institute for Biological Studies, 10010 North Torrey Pines Rd., La Jolla, CA 92037, USA, @kaymtye. Contributed Equally (Caroline Jia, Andrea Tran).

## Abstract

The “Pain Overlap Theory” (1) proposes that the experience of social pain overlaps with and amplifies the experience of physical pain by sharing parts of the same underlying processing systems (2–6). In humans, the insular cortex has been implicated in this overlap of physical and social pain, but a mechanistic link has not been made (2,4,5,7–9). To determine whether social pain can subsequently impact responses to nociceptive stimuli via convergent electrical signals (spikes) or convergent chemical signals (neuromodulators), we designed a novel Social Exclusion paradigm termed the Fear of Missing Out (FOMO) Task which facilitates a mechanistic investigation in mice. We found that socially-excluded mice display more severe responses to physical pain, disrupted valence encoding, and impaired neural representations of nociceptive stimuli. We performed a systematic biosensor panel and found that endocannabinoid and oxytocin signaling in the insular cortex have opposing responses during trials where mice were attending or not attending to the Social Exclusion events respectively, demonstrating distinct neuromodulatory substrates that underpin different states of Social Exclusion. We also found that intra-insular blockade of oxytocin signaling increased the response to physical pain following Social Exclusion. Together these findings suggest Social Exclusion effectively alters physical pain perception using neuromodulatory signaling in the insular cortex.

## MAIN TEXT

Evolutionarily, social bonds endow survival advantages, facilitating access to food resources, protection from predators, and higher rates of mating (10–14). To maintain these bonds, it has been suggested that as sociality emerged throughout evolution, social species repurposed neural systems initially designed to prevent physical harm to additionally safeguard against social separation by eliciting social pain (8,15).

Social pain, the emotional pain caused by aversive experiences with other individuals (1,16–18), is an umbrella term that includes experiences such as social isolation, social exclusion, social rejection, and social loss and can act as a signal that triggers a need state to help maintain social homeostasis (16,19–23). Social pain is a unique aversive experience, which is typically prolonged and multidimensional (5), and acts as an internal state that can modify future responses to environmental stimuli (24–30).

Multiple theories have proposed that the experience of social pain can modulate both emotional valence (31) and physical pain (1,2). Using fMRI studies, social and physical pain overlap has been implicated in multiple brain regions associated with physical pain, including the anterior insular cortex (aIC) (8). However, there are many possible mechanisms that could underlie an increase in Blood-Oxygen Level Dependent (BOLD) signal, and these possibilities include increased blood flow in the absence of changes in neural activity, or as an indication of neural dynamics. This led us to ask, how is social pain represented within the brain and how can its perception modulate physical pain?

There are multiple animal models that exist to generate an aversive social experience, including social rejection, social isolation, and social defeat stress (32–35). To establish a paradigm that would allow us to investigate specifically social exclusion, which would simultaneously also avoid physical injury and have a repeatable trial structure, we developed a novel Social Exclusion paradigm which we termed the Fear of Missing Out (FOMO) Task. During the FOMO Task, standard physical pain assays, and a valence discrimination task, we performed cellular-resolution microendoscopic calcium imaging, collected biosensor-mediated endogenous neuropeptidergic dynamics, and applied intra-insular pharmacology to understand how social pain can have a sustained influence on physical pain responses.

### Social Exclusion enhances behavioral responses to physical pain

We speculated that the Social Exclusion condition of the FOMO task may be aversive in multiple dimensions, including: (1) “frustration” at the inability to access resources, (2) “envy” from observing others obtain rewards, and (3) “social pain” associated with exclusion from the social group. While these specific subjective experiences do not have a ground truth readout, we predicted an escalation of responses with each additional dimension included in the Social Exclusion condition.

To represent each level of the gradient, the FOMO task contained three behavioral conditions (Fig. 1a). First, in all conditions, four co-housed mice were trained to associate a cue with delivery of a chocolate milkshake reward to a communal food basin. During each of the conditions, mice underwent 60 trials within a ∼1hr session. During the onset of each trial, a conditioned stimulus (CS) is presented and a switchable glass divider changed from opaque to transparent, allowing for visual access of the experimental side. Chocolate milkshake is then dispensed to the experimental side. All mice have demonstrated that they will pursue the reward, both individually and in a group setting (each mouse drinks 5 µl each trial during an individual setting, and collectively 15 µl during Social Exclusion).

**Fig. 1.**
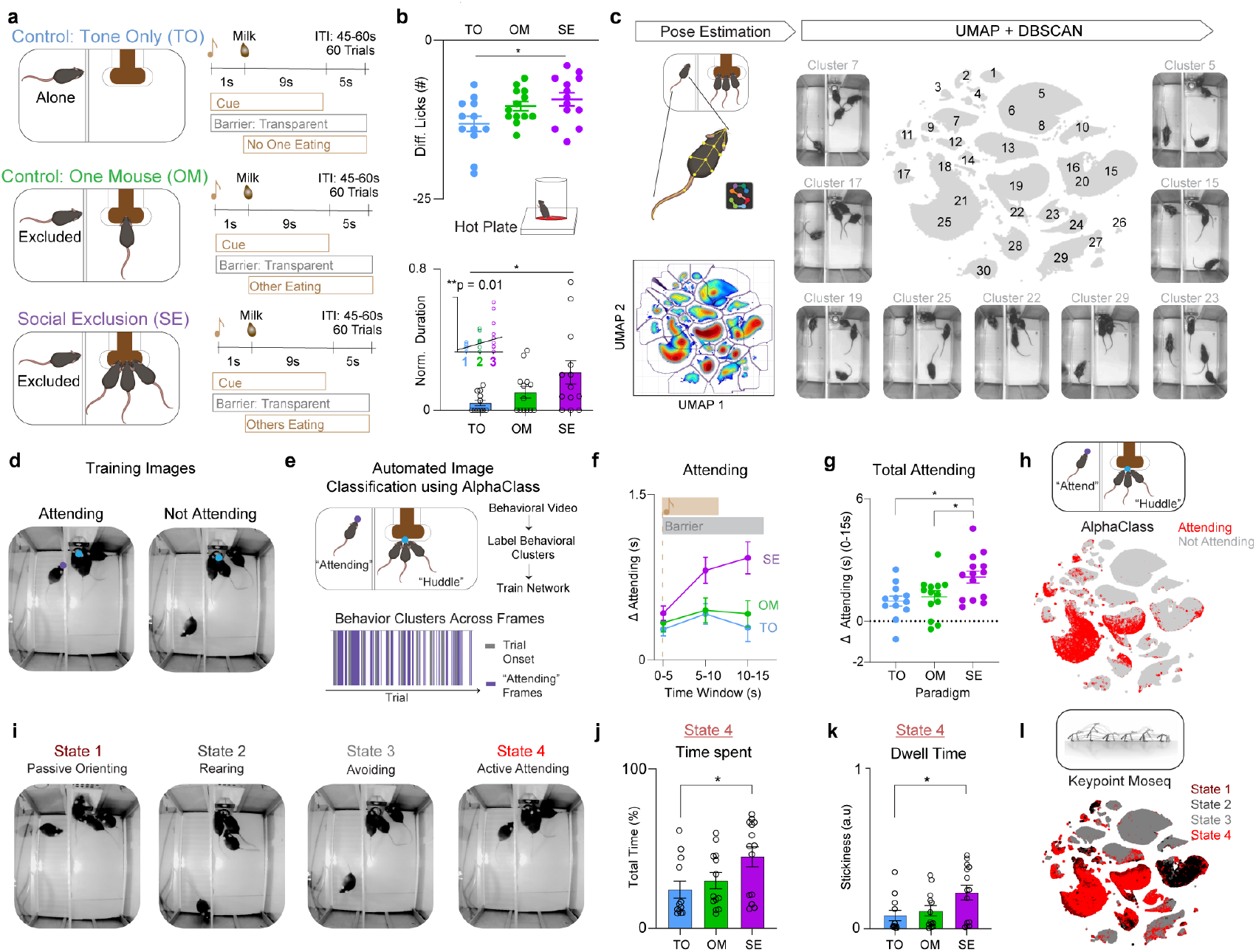
Social Exclusion enhances physical pain responses, reflected by increased licking on the hot plate. a, Experimental design for the Fear of Missing Out (FOMO) Task, which contains three behavioral conditions: Social Exclusion (SE), One Mouse (OM), and Tone Only (TO). Mice were presented with 60 trials for each session. At the beginning of each trial, the cue is on for 10s, followed by the delivery of chocolate milkshake 1s into the cue. The switchable glass divider between the social group and the excluded mouse turns transparent for 15s upon cue onset. Afterwards, we presented mice from all conditions with the hot plate assay to assess for changes in nociceptive thresholds. Each mouse only went through one of the three social conditions. b, Mice that underwent Social Exclusion displayed enhanced licking behavior while on the hot plate. Normalized Lick Duration, (which is normalized post-social condition licking / baseline licking) and Difference Lick Score (which is a difference score of post-social condition licking – baseline licking) was calculated with hot plate behaviors taken at a different baseline session. For Normalized Licks and Lick Duration, values closer to 1 indicate less change from baseline and values closer to 0 indicate greater change from baseline.(Top: n = 38, One-Way ANOVA, F_(2,35)_ = 3.85, *p = 0.03, Tukey’s multiple comparisons test, SE vs OM p = 0.72, SE vs TO *p = 0.028, OM vs TO p = 0.14, Bottom n = 38, One-Way ANOVA, F_(2,35)_ = 3.82, *p = 0.03, Tukey’s multiple comparisons test, SE vs OM p = 0.18, SE vs TO *p = 0.027, OM vs TO, p = 0.62, Inset: n = 38, Linear Regression R^2^ = 0.17 **p = 0.009). c, Behavioral analysis pipeline for the Social Exclusion condition. The subject mouse’s body parts were extracted first with SLEAP, then selected features of the excluded mouse were calculated. Next, we ran Uniform Manifold Approximation and Projection (UMAP) for dimensionality reduction and Density-Based Spatial Clustering of Applications with Noise (DBSCAN) for behavioral clustering. Displayed are example clusters generated through this pipeline and the UMAP for all frames. d, Example frames of “Attending” and “Not Attending” frames used to train AlphaClass, a new 2-D supervised machine learning algorithm for behavioral classification. e, “Attending” frames were used to train AlphaClass and all videos were run through the model to obtain “Attending” frames across the entire session. f, Quantification of time spent “Attending” during the 0-15s post cue period. ΔAttending is calculated based on the baseline period 5s before the trial starts. g, AUC taken from (f). Mice that undergo Social Exclusion spend more time engaging in “Attending” behaviors (n = 39, One-Way ANOVA, F_(2,36)_ = 5.77, *p = 0.01. Tukey’s multiple Comparisons Test: SE vs OM *p = 0.047, SE vs TO *p = 0.014, OM vs TO p = 0.85). h, AlphaClass classified frames are plotted onto the same UMAP template formed by DBSCAN to identify overlap between the different methods. Red color indicates “Attending” frames. Grey indicates “Not Attending” frames. i, An alternative clustering pipeline included used the behavioral features, combined with syllables from Keypoint-MoSeq (KPMS) into a hidden Markov Model (HMM) to discover four Hidden States. Example frames of four behavioral hidden states discovered through the HMM. State 4 is most similar to “Attending” behaviors identified through AlphaClass. j, Quantification of the time spent for State 4 among the three social conditions SE, OM, TO. Mice that underwent SE spend more time in State 4. (n = 39, One-Way ANOVA, F_(2,36)_ = 0.54, *p = 0.03, Tukey’s multiple comparisons test SE vs OM p = 0.15, SE vs TO *p = 0.035, OM vs TO p = 0.76). k, Mice that underwent SE stay in State 4 with a lower dwell time. Stickiness is calculated as a probability of remaining in the same state. (n = 39, One-Way ANOVA, F_(2,36)_ = 2.26, *p = 0.026, Tukey’s multiple Comparisons test, SE vs OM p = 0.09, SE vs TO *p = 0.03, OM vs TO p = 0.85). l, KPMS states are plotted onto the same UMAP clusters formed by DBSCAN. State 4 is overlapping with “Attending” clusters identified through AlphaClass. Error bars indicate s.e.m.

For the first dimension, inaccessible reward, we have a Tone Only (TO) condition, in which the CS tone is played, the switchable barrier becomes transparent, and the inaccessible reward is delivered at the trial onset. The One Mouse (OM) condition accounts for both the first and second dimension, the inability to access the reward and the distress of observing another obtain a reward (33,34). Lastly, to include all three dimensions, we have the Social Exclusion (SE) condition, in which mice are now excluded from a group of mice that are consuming the reward together, which is inaccessible to the subject. These conditions may all be aversive, but they allow us to differentiate between the contribution of the social group and/or the social agent relative to the denial of access to the chocolate milkshake.

After experiencing one of the three social paradigms (SE, OM, or TO), subjects were placed on a hot plate to assess nocifensive behaviors (behavioral responses to nociceptive stimuli (36)) (Fig. 1b). We found that Socially Excluded mice had a reduced relative change following the FOMO Task in comparison to the Tone Only group (Fig. 1b). Notably, we see a gradient of increasing nocifensive behaviors that correlates with an increasing number of dimensions of socioemotional challenges, from Tone Only with one, One Mouse with two, and Social Exclusion with three (Fig. 1b). Furthermore, this bias towards sustained, self-soothing behaviors (licking), compared to escape behaviors (jumping) suggests that Social Exclusion may selectively modify coping over escape affective behaviors (Extended Data Fig. 1a-c) (37–40).

### “Attending” behavior is robustly discoverable with multiple clustering methods

After validating that our novel Social Exclusion paradigm effectively modulates physical pain, we wanted to identify explicit behaviors performed during the paradigm that could help quantify the subjective experience of the excluded mouse. Previous work in the physical pain field has migrated to considering higher-order behavioral sequences to predict pain states (41), and we wondered if we could expand these efforts to dissect social distress (42).

To do so, we first extracted the subject’s behavioral outputs using pose-estimation software SLEAP (43) (Fig. 1c and Extended Data Fig. 2a). Next, we calculated various continuous and discrete behavioral features of the subject mouse, a subset of which included the distance and angle to the chocolate milkshake port, body velocity, acceleration, turning angle, and cage zone of the subject mouse (Extended Data Fig. 2b). We then leveraged dimensionality reduction and clustering methods to identify behaviorally relevant states from these high dimensional behavioral features using Uniform Manifold Approximation and Projection (UMAP) and clustered using Density-Based Spatial Clustering of Application with Noise (DBSCAN) (44) (Fig. 1c). Using this approach, we manually assigned clusters into two main subsets of behaviors, which we classified into either “Attending” or “Not Attending” behavioral clusters (Extended Data Fig. 2c). A behavioral cluster was labeled as “Attending” when the subject mouse was oriented towards, rearing, or climbing the switchable glass divider separating the subject from the reward delivery port and any other mice. All other behaviors were classified as “Not Attending”. Given that these labels were manually selected by human experimenters, we wanted to then use an unbiased approach with both supervised and unsupervised methods.

To validate the existence and relevance of these “Attending” behaviors, we developed a new supervised machine learning algorithm (AlphaClass) to determine if we could identify these behaviors across the entire session (Fig. 1d,e). AlphaClass is a behavioral segmentation method that classifies whether behaviors are occurring directly from single images using a convolutional neural network (CNN) (Extended Data Fig. 3). After training the network, we used AlphaClass to predict the presence of “Attending” behaviors on all frames in our behavioral videos. Using this method, we discovered that during each trial onset, there is a significant increase in “Attending” frames during Social Exclusion, compared to the OM and TO conditions (Fig. 1f,g). We found that if we mapped the frames that were classified as “Attending” back onto the UMAP-generated behavior embedding, AlphaClass successfully targets a subset of “Attending” clusters identified with DBSCAN (Fig. 1h). SE mice that exhibit more “Attending” behaviors also have subsequent enhanced nocifensive behavior (reduced latency to jump) on the hot plate (Extended Data Fig. 4b), as well as licks after formalin injection (Extended Data Fig. 4f) suggesting that not only can these “Attending” behaviors be used as a marker for social effort to reunite with the social group, but also these “Attending” behaviors correlate with multiple modalities of physical pain.

A disadvantage of these UMAP-DBSCAN and AlphaClass methods to identify “Attending” behaviors is that both methods assume static structure of data without incorporating the history of past behaviors, and we wanted to compare this to methods that were designed for kinematic sequences. Thus, to leverage the temporal component of our data, we also used an alternative unsupervised method merging Keypoint-Moseq (KPMS) (45) and a hidden Markov model (HMM) (25) to identify discrete, but prolonged behavioral states in an unbiased fashion. Within our dataset, KPMS identified 87 syllables (Extended Data Fig. 5b), which we merged with our behavioral features to obtain distinct prolonged behavioral states: Passively Orienting, Rearing, Avoiding, and Attending (Fig. 1i). We found that SE mice spend significantly more time and have a higher dwell time within State 4: “Attending” (Fig. 1j,k). When we map these specific four states back onto the UMAP template, we saw that they overlap with those defined as “Attending” by AlphaClass and DBSCAN (Fig. 1l and Extended Data Fig. 5a). This demonstrates that regardless of the clustering or machine learning algorithm, we can robustly discover an “Attending” social effort state that is elevated during Social Exclusion in methods-agnostic manner.

### Insular cortex cellular resolution neural responses to social and physical pain do not overlap

The insular cortex is a component of the “pain matrix” and interacts with higher order brain regions that process the emotional, affective, and cognitive aspects of pain (1,46–53). In rodents, the insular cortex has been shown to respond to both acute and chronic pain. Neuromodulators in the insular cortex, such as GABA, oxytocin (OXT), and endocannabinoids (eCBs), can induce antinociception and modify emotional states (46,54–58). In addition to the insular cortex’s role in pain behaviors, this region is also activated throughout social interactions and drives avoidance responses to social affective stimuli in rodents (59–61). The insular cortex is divided into the anterior and posterior regions, which have been shown to correspond with processing the affective and sensory components of pain respectively (47,59,62–64). This places the aIC in a unique position to integrate emotional and physical pain information. The aIC is poised to process and integrate aversive social information and utilize this information to regulate the impact of other sensory modalities, including that arising from physical stimuli evoking a nociceptive state.

The Pain Overlap Theory suggests that there would be convergence of social and physical pain signals in the same region (7), and there are multiple possibilities to explain the human fMRI co-activation for physical and social pain, which can occur either through cellular or population level convergence. One possibility is that the same cells code for cross-modal pain stimuli (electrical convergence). Alternatively, common neuromodulatory or neuropeptidergic signals could offer a biological mechanism for the Pain Overlap Theory (chemical convergence). Finally, it is also possible that there is no biological mechanism for convergence, and that the circuits that mediate physical and social pain do not actually converge but are just independently adjacent, recruiting a similar BOLD signal (7).

To uncover the mechanism of overlap between social and physical pain, we injected adeno-associated virus serotype 1 with a synapsin promoter for the genetically-encoded calcium indicator version 7f (AAV_1_syn-jGCaMP7f) in the aIC, implanted a GRIN lens to perform epifluorescent cellular-resolution calcium imaging (Fig. 2a) and recorded pan-neuronal aIC activity during the FOMO task (SE, OM, and TO) as well as during innocuous and nociceptive stimulus applications (Fig. 2a). For this study, we transitioned to more transient physical pain stimuli that could be repeatedly administered. To select for specific trials in which “Attending” behavior was highest and lowest, we used the top and bottom 15 trials with the most and least “Attending” frames respectively. We aligned neural activity to the onset of the cue during both “Attending” and “Not Attending” trials in the FOMO Task, and to the onset of pinprick on nociceptive trials (Fig. 2b). We co-registered neural responses to track the same neurons across all social conditions and physical pain stimuli (Fig. 2a).

**Fig. 2.**
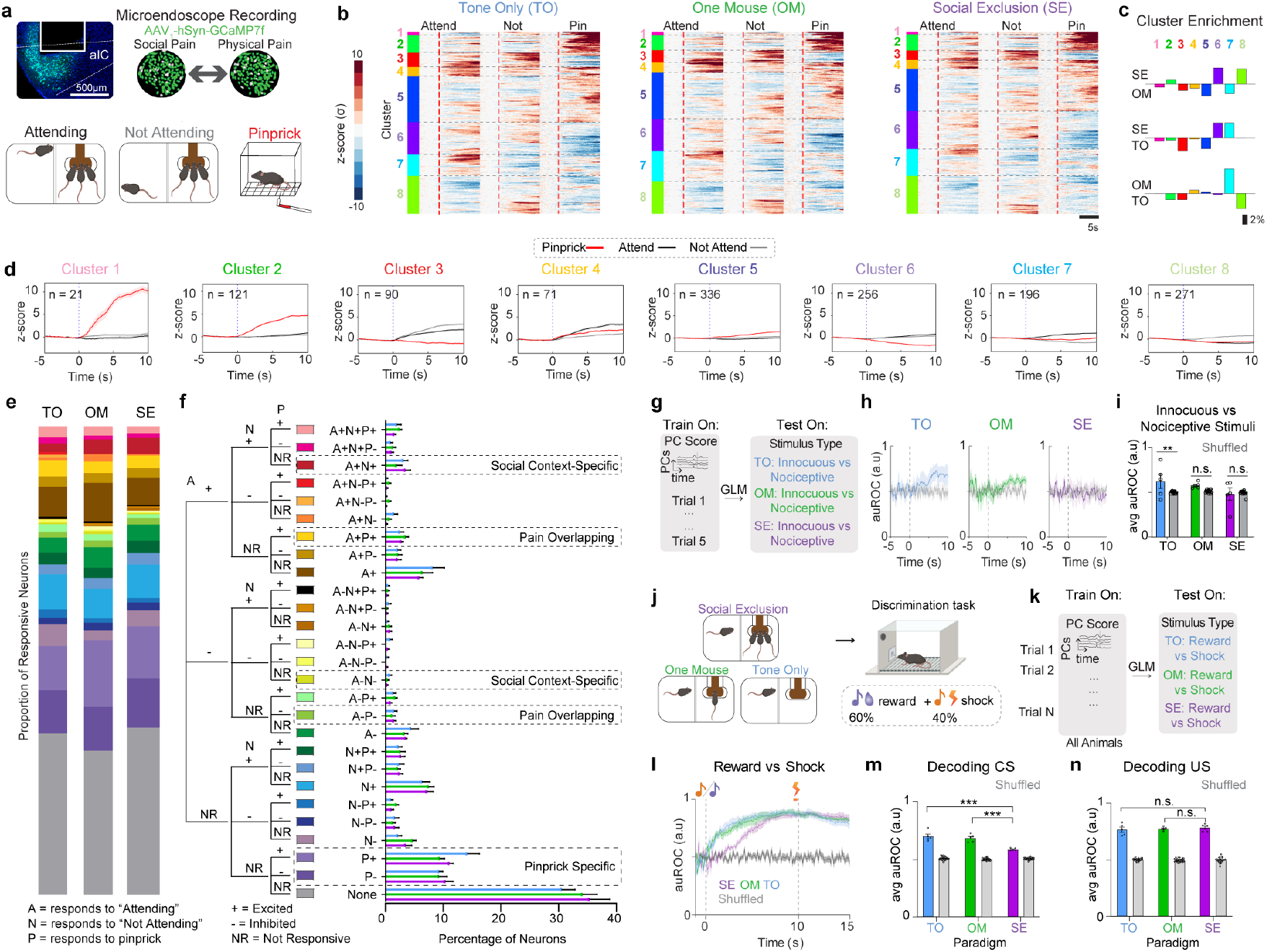
Social Exclusion impairs the ability to discriminate painful from innocuous stimuli or rewards. a, (Top) Example histological image of GCaMP7f expression in the aIC coronal section. Example output from cell co-registration to track the same neurons across different recording sessions. (Bottom) Example schematics of “Attending”, “Not Attending” and pinprick trials that were used for cellular resolution analyses. b, Neuron responses are organized using hierarchical clustering and plotted by condition. aIC ΔF/F_0_ responses to “Attending” and “Not Attending” trials are shown during SE, OM and TO, as well as during pinprick stimuli presented after each of the social conditions. Social trials are aligned to trial onset. At trial onset, the cue is played, and the switchable glass divider becomes transparent. Pain trials are aligned to stimulus application onset. Neurons are co-registered per social condition and physical pain stimulus presentation. Colors indicate clusters obtained using hierarchical clustering. c, Cluster enrichment calculated between social conditions. d, Population PSTHs to “Attending” (black), “Not Attending” (grey), and pinprick (red) trials for each cluster. e, Charts depicting the responsiveness of all neurons to “Attending”, “Not Attending” and pinprick trials. Colors in the bar graphs match the right graph. f, Dendrogram showing excitation and inhibition of neurons in response to the “Attending”, “Not Attending” and pinprick trials. For example, the first row of neurons is excited in response to “Attending” (A+), “Not Attending” (NA) and pinprick (P+) trials. Specific clusters that represent pain-overlapping neurons (responds to “Attending” and pinprick), social context specific neurons (responds to “Attending” and “Not Attending” trials), or pinprick specific neurons are indicated in dotted lines. There is no difference between conditions for any stimulus group. (Two-Way ANOVA, Main Effect across conditions p = 0.61, F_(2,15)_ = 0.52, Main Effect Across Responsiveness Types ***p < 0.0001, F_(4,60)_ = 160, Interaction p = 0.67, F_(52,390)_ = 0.9). g, Decoding schematic using a generalized linear model (GLM) to decode between different innocuous and nociceptive stimuli in each condition. Input data is PC transformed neural activity per animal during light touch, pressure, pinprick, and hot water trials. The number of PCs required to explain 90% of the variance was used in this analysis, and decoding was done using 5-fold cross validation (5 animals, five trials per animal per stimulus). Each stimulus was tested against a trial matched number of responses to other stimuli. This process was repeated three times to account for the total number of other stimuli trials. All trials were trial matched. h, Decoding performance for physical pain stimuli after SE, OM, and TO social conditions across all timepoints. i, Average decoding performance from 0-10s decoding between innocuous (light touch) vs nociceptive stimuli (pressure, pinprick, and hot water stimuli). Each data point represents the decoding performance per k-fold. Innocuous vs nociceptive stimuli are distinct after Tone Only, but not after One Mouse or Social Exclusion. (Two-Way ANOVA, Main Effect Shuffled Control. **p = 0.008, F_(1,54)_ = 7.582. Paradigm Effect: F_(2,54)_ = 4.65, *p = 0.01. Paradigm x Shuffled Interaction: F_(2,54)_ = 4.50. *p = 0.0156. Tukey’s multiple comparisons test, SE vs SE shuffled p = 0.88, OM vs OM shuffled p = 0.18, TO vs TO shuffled **p = 0.002). j, Schematic for Pavlovian discrimination behavioral task. Mice first went through one of the three social conditions: TO, OM, or SE, and then underwent a two-tone discrimination task. k, Decoding schematic using a GLM. Input data is PC transformed shock and reward trial data within each condition, with reward or shock trial labels. The number of PCs required to explain 90% of the variance was used in this analysis, and decoding was done using 5-fold cross validation (5 animals, 30 trials per animal per stimulus). l, Decoding performance at each timepoint between reward and shock trials using a GLM from 0-15s. Shock onset occurred at 9.8s for 200ms. CS onset occurred at 0s. Reward was delivered at 1s. Colors indicate behavioral condition, and the grey colored trace represents the shuffled control. m, AUC under the ROC curve from 0-5s post cue onset, to represent decoding performance of the conditioned stimulus. Mice that underwent SE have decreased decoding performance. Each data point represents the decoding performance per k-fold. (Two-Way ANOVA, Paradigm F_(2,84)_ = 54.51 ***p < 0.0001, Shuffled Control F_(1,84)_ = 1018 ***p < 0.0001, Paradigm x Shuffled Control F_(2,84)_ = 55.33 ***p < 0.0001. Tukey’s multiple comparisons test SE vs SE shuffled ***p < 0.0001. OM vs OM shuffled ***p < 0.0001, TO vs TO shuffled ***p < 0.0001. SE vs OM: ***p < 0.0001, SE vs TO: ***p < 0.0001, OM vs TO p = 0.37). n, AUC under the ROC curve from 10-15s post cue onset, to represent decoding performance of the unconditioned stimulus. Each data point represents the decoding performance per k-fold. There is no difference between the decoding performance for Social Exclusion compared to each of the controls. (Two-Way ANOVA, Shuffled Control F_(1,84)_ = 2302, ****p < 0.0001. Paradigm F_(2,84)_ = 0.45, p = 0.64, Paradigm vs Shuffled F_(2,84)_ = 1.11, p = 0.33). Error bars and solid shaded regions around the mean indicates s.e.m.

Previous work has suggested that specific ensembles of neurons can mediate state-dependent changes in behavior (65,66). To test if physical pain and Social Exclusion activate the same pain-responsive ensembles, we sorted these neural responses using hierarchical clustering and found clusters of neurons that differentially respond to social and physical stimuli (Fig. 2c,d and Extended Data Fig. 6e). We observed that all response profiles (Clusters 1-8) were seen in all conditions (SE, OM, TO) (Extended Data Fig. 6). To explore whether it is possible that Social Exclusion modulates physical pain through a separate and distinct pain-responsive population, we calculated the proportion of neurons that are excited by both social conditions and physical pain and found that there was no difference between the SE, OM, and TO groups (Fig. 2e,f). While this does not preclude the possibility that there still may be neurons that co-represent physical and social pain, we were unable to find significant evidence to support the biological model that physical and social pain converges onto the same neurons in aIC when using these stimuli.

### Ability of aIC neurons to decode painful stimuli is abolished after Social Exclusion

Since multiple groups have demonstrated that large scale cortical changes due to social behavior are spread across the entire population (67,68), we explored the possibility that pain could be represented via distributed coding across the aIC. To test for population level changes in physical pain encoding, we reduced the dimensionality of all population neural activity into principal components (PCs) using principal component analysis (PCA) (Fig. 2g). Using a support vector machine (SVM) trained on PC responses to innocuous and nociceptive stimuli, we found that the unique encoding of pinprick stimuli is abolished after both OM and SE conditions, but not the TO condition (Fig. 2h,i). We speculate this may underlie the heightened pain behaviors observed following Social Exclusion, mediated by enhanced sensory or emotional sensitivity to stimuli that are normally innocuous or only mildly painful. This demonstrated that the experience of Social Exclusion effectively modified physical pain encoding and led us to wonder if the changes in the representation of nociceptive stimuli within the aIC are due to changes in the representation of the sensory detection, or the affective, emotional component of physical pain.

### Social Exclusion disrupts valence representations within the aIC

Divergent ensembles within various cortical and subcortical regions have been demonstrated to play a differential role in mediating either the sensory-discriminative, cognitive, or affective components of pain (51,69–72). To test for changes in aIC encoding of the sensory or affective component of pain, mice underwent each of the social conditions and then performed a Pavlovian Discrimination task afterwards to evaluate changes in valence and sensory processing (Fig. 2j). We found that mice express no detectable single unit changes in either reward (Extended Data Fig. 8g) or shock trials (Extended Data Fig. 8h).

At the population level, we used a GLM to decode between reward and shock trials (Fig. 2k) and found that decoding performance is lower for Socially Excluded mice at the onset of the conditioned stimulus (CS) (Fig. 2l,m). However, this decoding performance change was not detected at the onset of the unconditioned stimulus (Fig. 2l,n), suggesting impaired discrimination of positive and negative associative memories with the reward- and punishment-predictive cues. This evidence supports the notion that Social Exclusion could change physical pain representation by modulating the affective emotional component of pain, while leaving the sensory component of pain intact.

### aIC encoding of social rank dynamically shifts based on FOMO Task condition

Next, we wanted to test if past social history could modulate responses to Social Exclusion by assessing the relationship between social rank and “Attending” behaviors. We hypothesized that social rank would differentially influence the amount of “Attending” behaviors but found no correlation between the number of “Attending” frames and the social rank of mice (Extended Data Fig. 9c). While we did not detect evidence that a single session of Social Exclusion impacts social rank decoding, it is possible with repeated Social Exclusion experiences that this would induce lasting changes to the group dynamics.

Although we did not observe a behavioral difference in “Attending” behaviors depending on social rank, we wondered if the representation of “Attending” and “Not Attending” social trials induced a different internal state depending on rank. Using a generalized linear model (GLM) to decode between social ranks based on aIC neural responses to all social trials, we find that rank information is decodable in all animals selectively during the Tone Only condition (Extended Data Fig. 9h). This is consistent with previous work that has shown social rank is encoded even when animals are alone (73). In contrast, during Social Exclusion, subordinate rank information is not decodable within the aIC, and during the One Mouse condition, intermediate rank decoding was impaired (Extended Data Fig. 9h). This suggests that the aIC dynamically shifts the encoding of rank information following conditions that may induce social pain, in a way that we speculate may be distinct for emotional primitives (74) of “envy” and “exclusion.”

### Oxytocinergic (OXT) and endocannabinoid (eCB) dynamics are recruited during Social Exclusion

After demonstrating impairments at the population level encoding of physical pain and valence after Social Exclusion, we searched for a molecular mechanism that could mediate this shift. Neuromodulation has been previously shown as a mechanism to bridge neural activity occurring at different timescales to guide behavior (18,66,75). It is possible that experiences of Social Exclusion trigger the release of neuromodulators, which could drive a negative motivational state or change neuronal excitability and thereby alter physical pain processing by recruiting more neurons to the pain-responsive ensemble.

Given that dorsal raphe nucleus (DRN) dopamine neurons play an important role in mediating an aversive state of social isolation (33) and that dopamine recruitment can be protective (76–80), we wondered if dopamine would, in addition to affecting the quantity of social interactions, also influence the quality of social interactions to preserve a level of social homeostasis (19). To test this, we expressed the GRAB_DA3h_ sensor (81), containing a G protein-coupled receptor (GPCR) with a circularly permutated fluorescent protein inserted into the third intracellular loop of the dopamine receptor, in the aIC and recorded neural activity during the FOMO task (Fig. 3a). We did not detect any significant differences in dopaminergic dynamics across the three conditions, SE, OM, and TO, between the “Attending” and “Not Attending” trials (Fig. 3a-d), nor were we able to decode trial type (“Attending” versus “Not Attending”) using a random forest classifier (Fig. 3e). Together, this suggested that dopamine in the aIC is not predictive of any of the experimental variables.

**Fig. 3.**
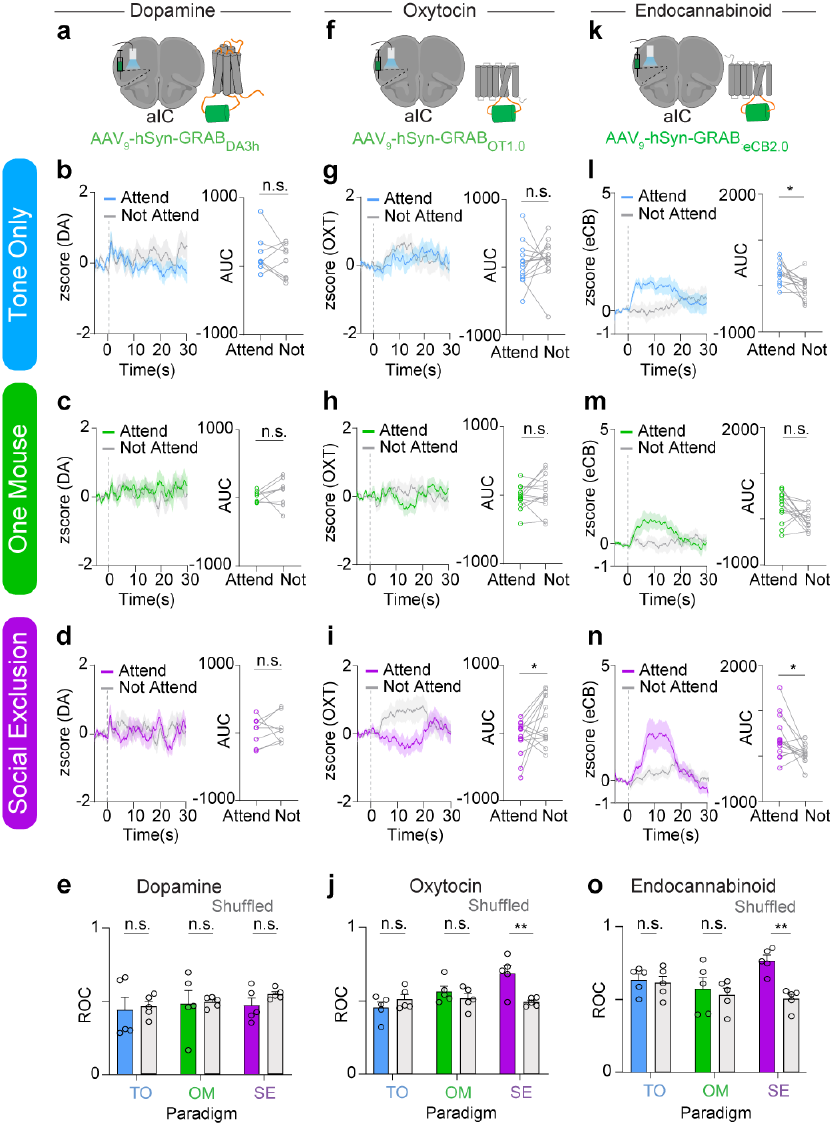
Genetically encoded fluorescent sensors reveal the role of oxytocin and endocannabinoid signaling dynamics during Social Exclusion. a, Fiber photometry and viral injection schematic using a genetically encoded dopamine sensor GRAB_DA3h_. b-d, Dopaminergic dynamics during Social Exclusion (SE), One Mouse (OM), and Tone Only (TO) conditions. The dotted line for each trace marks the onset of each trial. Each trial lasts 15s. 15 trials were taken to generate a trial average response, which was determined using the local thresholds per animal. The AUC was taken from 0-20s to account for sensor dynamics. (b) Tone Only responses (n = 8, two-tailed paired t-test, p = 0.35, t_7_ = 0.99). (c) One Mouse responses (n = 8, two-tailed paired t-test, p = 0.49, t_7_ = 0.72). (d) Social Exclusion responses (n = 8, two-tailed paired t-test, p = 0.68, t_7_ = 0.43). e, Decoding “Attending” and “Not Attending” trials within each social condition (SE, OM, and TO) using a random forest model and 5-fold cross validation. ROC values obtained from using dopaminergic dynamics to decode between “Attending” and “Not Attending” (Two-Way ANOVA, Main Effect across conditions p = 0.61, F_(2,24)_ = 0.50, Main Effect Data vs Shuffled p = 0.43, F_(1,24)_ = 0.64). f, Fiber photometry and viral injection schematic using a genetically encoded oxytocin sensor GRAB_OT1.0_. g-i, Oxytocinergic dynamics during Social Exclusion, One Mouse, and Tone Only paradigms, extracted from “Attending and “Not Attending” Trials. Baseline for z-score was - 5–0s prior to cue onset. Dotted lines at 0 represent cue onset. Area under curve was taken from 0-20s post cue. (g) Tone Only responses (n = 14, two-tailed paired t-test; p = 0.53, t_13_ = 0.64). (h) One Mouse responses (n = 13, two-tailed paired t-test; p = 0.29, t_12_ = 1.12). (i) Social Exclusion responses (n = 14, two-tailed paired t-test; *p = 0.039, t_13_ = 2.29). j, ROC values obtained from using oxytocinergic dynamics to decode between “Attending” and “Not Attending” (Two-Way ANOVA, Main Effect across conditions *p = 0.02, F_(2,24)_ = 4.4, Main Effect Data vs Shuffled *p = 0.047, F_(1,24)_ = 4.4, Interaction **p = 0.007, F_(2,24)_ = 6.1. Sidak’s Multiple Comparisons Test, SE vs SE shuffled **p = 0.003, OM vs OM shuffled p = 0.7, TO vs TO shuffled p = 0.6). k, Fiber photometry and viral injection schematic using genetically encoded endocannabinoid sensor GRAB_eCB2.0_. l-n, Endocannabinoid dynamics during Tone Only, One Mouse and Social Exclusion paradigms, extracted from “Attending” and “Not Attending” Trials. Area under curve was taken from 0-15s post cue. l, Tone Only responses (n = 13, two-tailed paired t-test; *p = 0.01, t_13_ = 2.9). m, One Mouse responses (n = 14, two-tailed paired t-test, p = 0.05, t_14_ = 2.2). n, Social Exclusion responses (n = 14, two-tailed paired t-test, *p = 0.02, t_13_ = 2.6). o, ROC values obtained from using endocannabinoid dynamics to decode between “Attending” and “Not Attending” (Two-Way ANOVA, Main Effect across conditions p = 0.2, F_(2,24)_ = 1.7, Main Effect Data vs Shuffled *p=0.01, F_(1,24)_ = 7.6, Sidak’s Multiple Comparisons Test, SE vs SE shuffled **p = 0.002, OM vs OM shuffled p = 0.9, TO vs TO shuffled p = 0.99). Error bars and solid shaded regions around the mean indicates s.e.m.

The oxytocinergic system is well known to be a key regulator of social bonds (46,76,82–89), and could contribute to a neuropeptidergic mechanism that protects against Social Exclusion. Furthermore, oxytocin has also been shown to have widespread analgesic properties in the central nervous system (46,84,90,91). Thus, it is possible that oxytocin could modulate both Social Exclusion and physical pain and act as a neural substrate that bridges the distinct timescales of Social Exclusion and physical pain. To test if oxytocinergic signaling occurs during Social Exclusion, we expressed the GRAB_OT1.0_ biosensor (92), which has a circularly permutated GFP, inserted into the third intracellular loop of the oxytocin receptor, in the aIC and recorded oxytocinergic activity using bulk fluorescence (Fig. 3f). No detectable difference was observed during Tone Only and One Mouse conditions in OXT signaling during “Attending” and “Not Attending” trials (Fig. 3g,h). However, in the Social Exclusion condition, oxytocin signaling is increased during “Not Attending” trials, compared to “Attending” trials (Fig. 3i), suggesting a protective effect of oxytocin that is selective for Social Exclusion. Further, the same random forest classifier that returned null results for the dopamine sensor (Fig. 3e) showed significant decoding of “Attending” and “Not Attending” trials based on the GRAB_OT1.0_ readout in the Social Exclusion condition, but not the One Mouse or Tone Only conditions (Fig. 3j), further supporting the notion that oxytocin is reduced during “Attending” trials, in a manner specific to being excluded from a social group.

The endocannabinoid (eCB) system has been implicated in the regulation of physical pain behaviors (93,93–96), and has also been shown to induce antinociception, suppress nociceptive behaviors, and alter affective behaviors (96–99). eCBs have also been implicated in social reward and social interest (100–103), and therefore, could also play a role during Social Exclusion. Finally, the eCB system has long been recognized to be important for modulating food intake (104,105) and eCBs in the aIC have been implicated in water intake (106).

To test the role of endocannabinoid signaling during Social Exclusion, we expressed the GRAB_eCB2.0_ biosensor (107), containing a circular-permutated EGFP and the human CB1 cannabinoid receptor, in the aIC using viral transduction and recorded endocannabinoid activity using bulk fluorescence (Fig. 3k). We discovered that eCB signal was also present during “Attending” behaviors during the Tone Only condition as well and experiencing omission of the chocolate milkshake reward (Fig 3l-n) suggesting that it is possible that the eCB signal integrates food reward and social information. Even so, when we used the same random forest classifier to decode “Attending” versus “Not Attending” trials from the GRAB_eCB2.0_ biosensor, we were only able to get significant decoding in the Social Exclusion condition, and not in the Tone Only nor One Mouse conditions (Fig. 3o).

Taken together, we used three different biosensors in the aIC to measure dopaminergic, oxytocinergic and endocannabinoid dynamics during each condition of the FOMO Task, and found that bulk fluorescence from both oxytocin (Fig. 3j) and endocannabinoid (Fig. 3o), but not dopamine (Fig. 3e), biosensors could be used to selectively decode trial types in the Social Exclusion condition, but not in the One Mouse nor Tone Only conditions. Notably, the signals measured by the oxytocin (Fig. 3f-j) and endocannabinoid (Fig. 3k-n) biosensors had opposing directionality with the GRAB_OT1.0_ readout elevated during “Not Attending” trials and the GRAB_eCB2.0_ fluorescence elevated during “Attending” trials (Fig. 3f-o). Despite this opposing signal directionality, both biosensors allowed trial types to be significantly decodable selectively during the Social Exclusion condition (Fig. 3j and 3p). To our knowledge, this represents the first neural substrate specific to social exclusion, and this neuropeptidergic signature points to a vast space capable of multiplexing to mediate complex socioemotional states.

### Blockage of Oxytocinergic (OXT) signaling enhances physical pain responses after Social Exclusion

Given the analgesic properties of oxytocin and endocannabinoids (46,96,108), we speculated that the blockade of oxytocinergic signaling could promote active coping in response to painful stimuli, including “Attending” behavior in the FOMO Task or licking on the hot plate. Previously, it has been shown that direct injection of oxytocin into the aIC has anti-nociceptive effects (46). To first test the causal role of oxytocin or endocannabinoids, we injected an oxytocin receptor antagonist (OXTRA) L-368,899 or cannabinoid receptor 1 agonist (CB1RA) WIN55,212-2 into the aIC and exposed mice to Social Exclusion followed by the hot plate (Fig. 4a). To better visualize the specific granular behaviors modified by these pharmacological agents, we mapped behavioral responses back onto the same UMAP in Figure 1 to reveal the detailed behavioral profiles produced by CB1 receptor agonism and OXT receptor antagonism relative to their respective vehicle controls (Fig. 4b-d). Here, we focus on behaviors related to Attending to the other side of the chamber, but this detailed quantitative readout can be used to inform future research on other aspects of behavior.

**Fig. 4.**
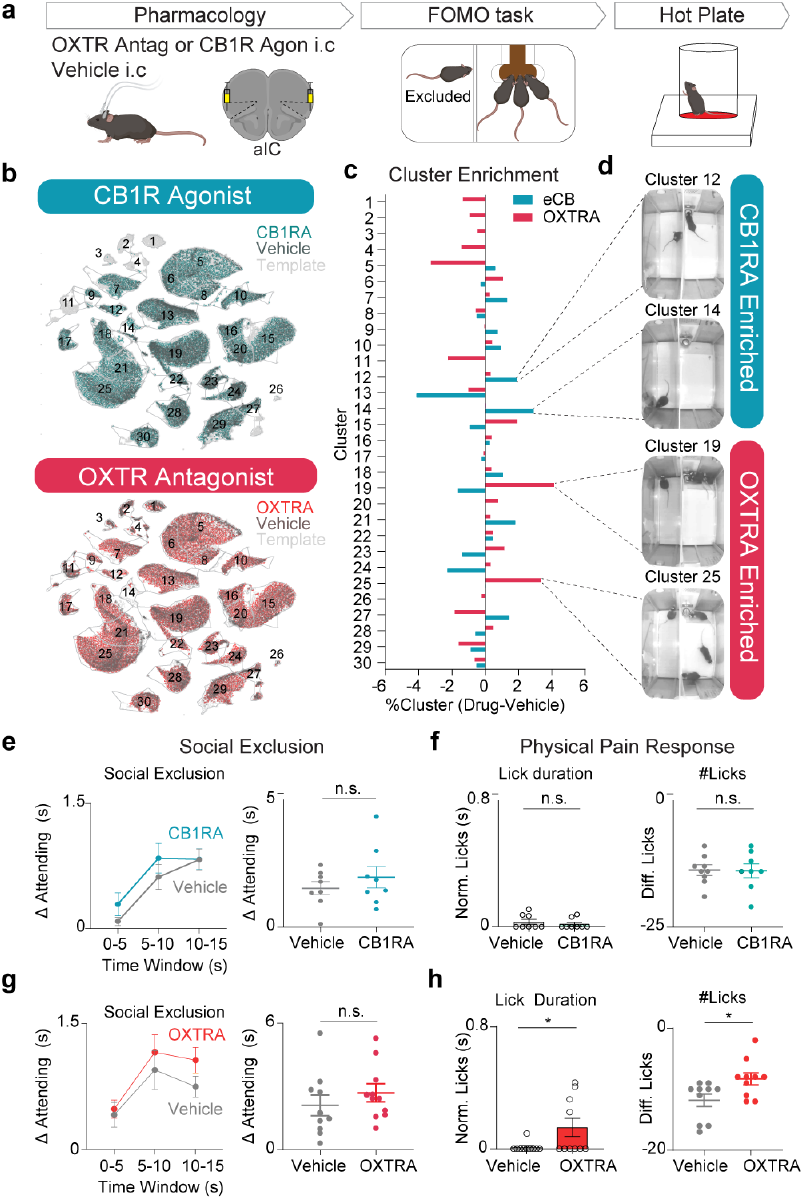
Genetically encoded fluorescent sensors reveal the role of oxytocin and endocannabinoid signaling dynamics during Social Exclusion. a, Schematic of intracranial injection of L-368,899 oxytocin receptor antagonist and WIN55,212-2 cannabinoid receptor agonist into the aIC. Mice are run 10 minutes post-injection through Social Exclusion, and subsequently, the hot plate. b, UMAP plots depicting the amount of frames that belong to each cluster in conditions: (top) OXTRA and Vehicle or (bottom) CB1RA and Vehicle. c, Enrichment of behavioral frames for each cluster. Bars are obtained by %Drug frames in cluster - %Vehicle frames in cluster. d, Example frames of clusters that are enriched after OXTRA or CB1RA administration. e, “Attending” behavior between CB1RA and vehicle controls. There is no difference in “Attending” behavior. Normalized time spent “Attending” was calculated throughout the entire 15s trial. (n = 8 vehicle, n = 8 CB1RA, two-tailed unpaired t-test, p = 0.39, t_14_ = 0.89. f, (Left) Normalized lick duration during CB1RA pharmacology experiments. (n = 16, two-tailed unpaired t-test, p = 0.48, t_14_ = 0.72) (Right) Hot plate difference lick score between CB1RA and Vehicle controls. (n = 16, two-tailed unpaired t-test p = 0.94, t_14_ = 0.07). g, “Attending” behavior between OXTRA and vehicle controls. There is no difference in “Attending” behavior. Normalized time spent “Attending” was calculated throughout the entire 15 second trial. (n = 10 vehicle, n = 10 OXTRA, two-tailed unpaired t-test, p = 0.37, t_18_ = 0.92). h, (Left) Normalized lick duration during OXTRA pharmacology experiments after Social Exclusion. Lick duration is increased with OXTRA injection. Oxytocin antagonism increases social exclusion-induced pain (two-tailed unpaired t-test *p = 0.045, t_18_ = 2.16) (Right) hot plate difference lick score between OXTRA and vehicle controls. Normalized licks are increased with OXTRA injection. (n = 20, two-tailed unpaired t-test, *p = 0.023, t_18_ = 2.49). Error bars and solid shaded regions around the mean indicates s.e.m.

Then we used AlphaClass to quantify the total “Attending” behavior during Social Exclusion and found that neither of the manipulations changed total “Attending” behavior during Social Exclusion (Fig. 4e,g). However, we did find that OXTRA mice exhibit an increase in affective nocifensive behaviors during hot plate (licking; Fig. 4h), providing evidence that OXT is protective against increased affective physical pain behaviors following Social Exclusion.

Collectively, our work provides the framework for a novel paradigm to induce Social Exclusion and the first evidence for a specific neural mechanism for the overlap between Social Exclusion and physical pain. Our findings support the “Pain Overlap Theory” and suggest that the oxytocinergic system is a key mediator of Social Exclusion within the aIC, which can successfully act as a bridge between different types of aversive experiences by modulating negative affect. Furthermore, this advances our general understanding of neural strategies that are employed within emotional pain processing systems within the brain and paves the way for targeted interventions aimed at alleviating the distress associated with social disconnection and pain dysregulation.

## Acknowledgements

This work was supported by the Machine Shop of the Salk Institute and Bryan Nielsen. This preprint used the zHenriquesLab template for Overleaf and diagrams from Biorender. Funding: Salk Institute for Biological Sciences (KMT) HHMI (KMT) Kavli Foundation (KMT) Clayton Foundation (KMT) Dolby Family Fund (KMT) NIMH R01-MH115920 (KMT) NIMH R37-MH102441 (KMT) NICCIH Pioneer Award DP1-AT009925 (KMT) Salk Women in Science (CJ) NIMH 1RF1MH132653-01 (TDP) NIDA R00DA055111 (HL) Author Contributions: Conceptualization: CJ, KMT Stereotaxic Surgeries: CJ, AT Behavioral Experiments: CJ, AT, FA, NT, ES, NNN Fiber Photometry and Calcium Imaging Experiments: CJ, AT, FA, AN SLEAP automated pose tracking analysis: CJ, MGC, AT, FA, NT, ES, NNN Keypoint-Moseq Analysis: CJ, ES Histological Verification: CJ, AT, FA, ES, NT Code Scripts: CRL, RRP, KB, JD, LRK Intellectual contributions: FT, KB, RW, MKB, TDP, HL AlphaClass architecture: AB Data analysis: CJ, AT, ES, NT, FA, AN Data preparation and sharing: CJ, AT, RW, LRK Materials: YL Writing – original draft: CJ, KMT Writing – review and editing: all authors contributed to editing the manuscript

## Data and Materials Availability

All experimental data are available in the main text or supplementary material. Raw neural recordings and simultaneously recorded behavioral outputs are available on DANDI. AlphaClass package is available on the Tyelab GitHub.

## Supplementary Materials

Materials and Methods

Figs. S1 to S10

Videos S1 to S2

## MATERIAL AND METHODS

### Animals and housing

Adult, male C57/BL6J mice (8-12 weeks) from Jackson Laboratory were used for all experiments described. Mice were housed 4 mice/cage on a 12-h reverse light/dark cycle with ad libitum access to food and water unless otherwise specified. All experiments were conducted during the dark cycle phase and performed either in low white light or red light. All experimental procedures were carried out in accordance with NIH guidelines and approval of the Salk Institutional Animal Care and Use Committee.

### Stereotaxic surgeries

All surgeries were conducted under aseptic conditions. Briefly, mice were anesthetized with an isoflurane/oxygen mixture (4-5% for induction, 1-2% for maintenance) and placed in a stereotaxic head frame (David Kopf Instruments, Tujunga, CA, USA). A heating pad was placed under the mice to maintain body temperature, and Sterile Lubricant Eye Ointment (Stye INSIGHT Pharmaceuticals Corp. Langhorne, PA) was applied to the eyes to prevent drying. The incision area was shaved, and the skin was cleaned with alternating washes of 70% alcohol and betadine. A subcutaneous injection of lidocaine (0.5%) was placed at the incision site for 3-5 minutes prior to surgery. An incision was made along the midline to expose the skull, and a dental drill was used to perform a craniotomy. During all surgeries, animals were injected subcutaneously with 1 mL of Ringer’s solution, Buprenorphine (1 mg/kg), and Meloxicam (5 mg/kg). For recovery animals, were placed in a clean cage on a heating pad. Animals were given >28 days for recovery before starting behavioral paradigms.

All stereotaxic coordinates were measured relative to bregma and the top of the skull. Injections of viral vectors were performed using glass pipettes (Drummond Scientific) pulled to a 100-200 µm tip diameter with a pipette puller (Narishige PC-10, Amityville, NY, USA). Pipettes were either attached to 10 µl microsyringes (Hamilton Microlitre 701, Hamilton Co., Reno, NV, USA) with a microsyringe pump (UNP3; WPI, Worcester, MA, USA) and digital controller (Micro4; WPI, Worcester, MA, USA), or to the Nanoject III Programmable Nanoliter Injector (Drummond Scientific, Broomall, PA, USA) with digital controller (Drummond Scientific, Broomall, PA, USA). For each injection, micropipettes were slowly lowered to the target site and viral vectors were delivered at a rate of 1.0 nL per second. After the injection was completed, micropipettes were raised to 0.1 mm above the injection site and held for 10 minutes. After 10 mins, the pipette was slowly withdrawn, and the skin incision was closed with nylon sutures.

To perform calcium imaging recordings, mice underwent surgery as described previously. For non-specific aIC recordings, 200 nl of AAV_1_ -Syn-jGCaMP7f-WPRE (Addgene), encoding GCaMP7f, was injected into the aIC (AP: +1.9 mm, ML: +2.85 mm, DV: −3.4 mm) and a 0.6 mm diameter by 7.3 mm length gradient refractive index lens with integrated baseplate (GRIN lens, Inscopix) was slowly lowered above the aIC (AP: +1.9 mm, ML: +2.85, DV: −3.15 mm). No tissue was aspirated. All lens implants were secured to the skull with a thin layer of adhesive cement (C&B Metabond, Parkell), followed by black cranioplastic cement (Ortho-Jet, Lang). The implant was allowed to completely dry before closure of the incision with nylon sutures.

To record oxytocinergic, dopaminergic and endocannabinoid dynamics within the aIC, mice underwent surgery as described previously. An AAV_9_ carrying either the dopamine, oxytocin, or endocannabinoid fluorescent sensor (AAV_9_ -hSyn-GRAB_DA3h_, AAV_9_ -hSyn-GRAB_OT1.0_, AAV_9_ -hSyn-GRAB_eCB2.0_) was injected (AP: +1.9 mm, ML: +2.85 mm, DV: −3.4 mm) and a 400 µm diameter optical fiber was implanted into the aIC. All fiber implants were secured to the skull with a thin layer of adhesive cement followed by black cranioplastic cement. The implant was allowed to completely dry before closure of the incision with nylon sutures. Behavioral experiments were conducted 4-7 weeks after surgery.

To perform intracranial pharmacology within the aIC, mice underwent surgery as described previously. Bilateral cannulas (Protech Technologies) were implanted 1 mm above the aIC (AP: +1.9 mm, ML: +2.85 mm, DV: −2.4 mm). The incision was closed with nylon sutures. Behavioral experiments were conducted 2-3 weeks after surgery. During experiments, an internal cannula with 1 mm projection was used to deliver drug into the brain.

### Behavioral assays

Prior to behavioral testing, mice were habituated for one week to all experimenters to reduce stress during experiments.

### Food Restriction

Subjects were fed ad libitum until initiation of experiments requiring food restriction (reward conditioning and FOMO tasks). Prior to these experiments, subjects were weighed to the nearest gram to attain a baseline for free-feeding body weight. During experiments requiring restriction, subjects were weighed daily and food-restricted to a limit of 85% of this baseline weight. Restriction was attained through limiting daily access to food pellets, providing approximately 2.0g/animal per day. Animals had free access to water throughout.

### Reward conditioning

Food-restricted mice were conditioned in sound-proof boxes (MedAssociates, St Albans, VT) for five days. Each box contained a modular test cage assembled with a 3D-printed reward port, a speaker, a red LED indicator light, and 2 house lights. The first trial began after a 245s habituation period. The CS consisted of a pure 3.5kHz tone cue, which ended 400ms after a port entry infrared beam break by the mouse was detected. One second after CS onset, 5 µl of chocolate milkshake, Ensure™, was delivered. The first chocolate milkshake delivery was given freely but subsequent chocolate milkshake deliveries only occurred if the mouse had entered the port after the CS onset. Inter-trial time intervals varied between 40-60s. Each conditioning session consisted of 120 trials. Mice were considered successfully trained if they reached a 70% probability of port entry during the CS.

Three days into reward conditioning, the mice were placed in the same sound-proof boxes (MedAssociates, St Albans, VT) with a switchable glass divider connected through an IoT relay. Each mouse was individually placed on the side of the wall opposite of the reward port. Each habituation session consisted of 60 trials, during which no tones or reward were delivered. The purpose of this session was to habituate mice to the sound of the switchable glass divider.

### FOMO Task: Social Exclusion, One Mouse, and Tone Only conditions

Reward conditioned mice were placed in sound-proof boxes (MedAssociates, St Albans, VT), which contained a switchable glass divider, connected through an IoT relay, which divided the arena in half and was used to enforce a trial structure. Socially excluded mice were placed on one side of the wall, opposite of the chocolate milkshake-dispensing port and three cagemates. Similar to the reward conditioning trials, at the beginning of each trial, a 3.5kHz tone cue was presented and the dividing barrier turned transparent. This allowed the socially excluded mouse to visualize the other half of the box. One second after each trial onset, 15µL of chocolate milkshake was delivered to the port to the cagemates. The 3.5kHz tone was turned off after 10s, and the switchable glass wall transitioned back to opaque after 15s. Inter-trial time intervals varied between 40-60 seconds with an initial 245s delay for habituation preceding the first CS onset. Each session consisted of 60 trials.

For the One Mouse control, only 1 cagemate is on the other side and 5 uL of chocolate milkshake is delivered to the port during each trial. All other experimental methods are consistent with the Social Exclusion condition. For the Tone Only control, no cagemates are present on the other side, and 5 uL of chocolate milkshake is delivered to the port during each trial.

### Hot Plate

To measure thermal sensitivity after the Social Exclusion, One Mouse, or Tone Only behavioral conditions, mice were placed inside a cylindrical, transparent Plexiglass chamber (Diameter = 11 cm, Height = 15 cm) on a hot plate (54°C, IITC Life Science) for 60s. The latency and duration of each behavioral response (hind paw shake, lick or jump) was manually recorded using Behavioral Observation Research Interactive Software (BORIS) scoring software (105). To ensure unbiased evaluations, the manual scoring on BORIS was done without the knowledge of the experimental conditions. Because individual mice have been known to have variable responses on hot plate depending non-nociceptive factors (weight, age, activity, habituation and repeated testing), we calculated baseline values for each mouse’s baseline pain response (106,107). Baseline values were determined for each mouse’s baseline pain responses, based on a baseline hot plate session. Normalized Lick Duration for hot plate was calculated Paradigm / Baseline. Difference Lick Score was calculated using Paradigm-Baseline.

### Formalin Assay

To measure inflammatory responses after Social Exclusion, One Mouse, or Tone Only behavioral conditions, animals were injected in the right hindpaw with 1% formalin solution and placed in a rectangular, transparent Plexiglass chamber for 1 hour. The number of licks and duration of lick bouts was quantified using AlphaClass, which was trained on a model using 3700 frames.

### Innocuous and Nociceptive Stimulus Application

To measure time locked responses to various mechanical and thermal stimuli, mice were habituated to a clear plexiglass container (Animal enclosure, IITC433) on top of a mesh stand (IITC 410). On test day, after running through one of the three social paradigms, mice were placed within the plexiglass enclosure, and five rounds of four stimuli (0.16g Von Frey (IITC), 2.0g Von Frey, Pinprick using a 25g needle, 55°C Water Droplet) were applied. Each stimulus was applied in an alternating fashion to each hindpaw with one minute intertrial intervals and the neural response during each stimulus application was recorded and analyzed.

### Tube Dominance Test

The tube dominance test was used to assay the social rank of each mouse relative to their cagemates (108). Mice were individually trained to walk through a clear Plexiglass tube (30 cm length, 3.2 cm inner diameter) over ∼2days, until they were comfortably walking across without resistance. To test for rank, all mice in each cage were tested in a round robin design in a randomized order. For each pair, mice were released at opposite ends of the tube simultaneously, so that they met faceto-face in the center of the tube. The mouse that either backed out itself or was pushed out from the end where it was released was designated as “loser/subordinate” whereas the other mouse was designated as “winner/dominant.” Social ranks obtained with the tube test were considered stable when obtaining the same results for four or more days in a row. An animal’s “social rank” was measured by the proportion of “wins” across all contests from all days of testing.

### Pharmacological manipulation

OXT Receptor Antagonist (L-368,899 hydrochloride, Tocris Cat. No. 2631) was dissolved in sterile phosphate buffered saline (PBS) (50 µg/µl), aliquoted, and then stored in −20 °C. Drug was freshly dissolved each day. Endocannabinoid Receptor Agonist (WIN55212-2, Cayman Chemical No. 10009023) was dissolved in a 1:1:1:17 mixture of Ethanol, DMSO, Kolliphor, PBS, and then stored in −20°C.

For both drugs, 10 minutes prior to the start of the behavioral assay 0.2 µl was infused bilaterally into the insular cortex via dual internal infusion needles connected to a 10 µl microsyringe. The flow rate was kept to 100 nl per min and was regulated by a syringe pump (Harvard Apparatus, MA). Infusion needles were withdrawn 2 minutes after the infusion was complete. Each mouse only received one injection and each condition (OXTRA/CB1RA or Vehicle) utilized separate cohorts.

### Freely moving behavior, discrimination task

To test valence discrimination, mice underwent a Pavlovian discrimination assay. Associative reward training was performed as described above in reward conditioning. For shock acquisition, the mice were conditioned using behavioral hardware boxes (MedAssociates, St Albans, VT) placed in custom made sound attenuating chambers. Each box contained a modular test cage with an electric floor grid and a speaker. Videos of the mice were acquired during all sessions. A period of acclimation lasting four minutes preceded the presentation of the first tone. During shock conditioning, mice were presented with a 20kHz tone followed by a 0.1s long 0.7mA shock 9.8s later. Inter-trial intervals varied from 40-60s with an initial 10s delay before the first cue onset. Shock-tone association training consisted of 60 trials within one session.

One day after chocolate milkshake reward and shock conditioning, mice completed the Social Exclusion, One Mouse, or Tone Only social paradigm, and immediately underwent the two-cue Pavlovian discrimination task. During each trial, mice were presented with either a 3.5kHz tone (associated with chocolate milkshake reward) or a 20kHz tone (associated with shock) using the same trial structures as described above. Reward (56) and shock (34) trials were presented in pseudorandom order. Intertrial intervals were varied from 40-60s with an initial 4-minute delay for habituation preceding the first CS onset. An LED light was toggled on and off for the period of the tone and used to sync videos to trial start for data processing. Videos were recorded to monitor animal behavior during all sessions. Neural recordings using the Inscopix system described below were also obtained during each session. Each session consisted of 90 trials and the protocol was repeated for each of the Social Exclusion, One Mouse, and Tone Only social conditions for each mouse.

Freezing and dashing behavior were scored in Matlab by setting a threshold for velocity and acceleration for movement. Frames below this threshold were scored as freezing behavior.

### SLEAP automated pose tracking analysis

To automatically detect mouse social interaction behavior, Social LEAP Estimates Animal Poses (SLEAP), version 1.3.0 (43), was used to estimate animal poses in behavioral videos. Behavioral videos were taken using a video camera (Arducam 1080P, 30 fps). A training data set was labeled using a 9-point skeleton on the mouse (nose, right ear, left ear, torso, left forepaw, right forepaw, left hindpaw, right hindpaw, and tail base). All annotators were instructed to annotate the ears within the middle of each ear, the nose at the tip of the mouse nose, and the tail base where the tail began. Any body parts that were not clearly visible were empirically estimated to their most likely location given the mouse’s position. This data set was used to train a top-down model with 2955 frames, with −180° to 180° augmentation because videos were taken from above. All frames were visually confirmed for correct tracking of identity and manually corrected into a consistent track for the excluded animal.

### Low Dimensional Embedding of Behavioral Features and Unsupervised Clustering

We performed unsupervised classification of mice behaviors during the FOMO task. Each mouse was video recorded performing the social task and videos were subsequently labeled using automated pose estimation algorithms SLEAP as described above. To identify specific behavioral motifs, we extracted various behavioral features of the excluded mouse from SLEAP-predicted coordinates of labeled body parts. These coordinates were smoothed using the smoothdata function in MATLAB and interpolated to account for any missing keypoints. We extracted both continuous and discrete features: including the distance to port, angle to port, body velocity, body acceleration, orientation of head to body, orientation of head to tail, turning angle, 1second tortuosity, 5s tortuosity, distance of nose to tail, and change in distance of nose to tail, zone of excluded mouse, huddle quality (1, 2, 3), touching barrier, oriented to port, oriented and touching barrier. All continuous features were z-scored. Tortuosity is calculated as the ratio between the length of the path and the distance between the beginning and endpoints of the path. This feature can function as a proxy of the mouse’s path and can capture behavioral motifs in which the mouse takes longer or sharper turns. Each of these features were extracted for the 15s following each cue onset during the FOMO task. To identify distinct behavioral motifs mice exhibited during social interaction, we applied Uniform Manifold Approximation and Projection (UMAP) onto these behavioral features and obtained unsupervised clusters using DBSCAN. Clusters were manually annotated as an “Attending” or “Not Attending” cluster based on visualization of each cluster by generating a video from the frames included in that cluster.

### Keypoint Moseq (KPMS)

To perform unsupervised classification of mouse behaviors during the Social Exclusion, One Mouse, and Tone Only social paradigms using keypoint-MoSeq (45) (KPMS), which is built from an auto-regressive hidden Markov model (AR-HMM). The output of SLEAP, raw “keypoint” data for each frame, was used as training data for KPMS. The initial AR-HMM model was run through 200 iterations on a sample of 20 collected videos using a kappa value of 10e12. The full model was then run using a kappa value of 10e9 for 500 iterations. The final KPMS output was 87 “syllables” representing behavioral motifs in the data.

Together, the KPMS syllables were combined with seven other features: five continuous features: (distance of nose to port, angle of nose to port, body velocity, turning angle) and two discrete features (zone, oriented to port). KPMS syllables and seven features were run through a second HMM (adapted code from Vidaurre et al., 2017 (109)) to further consolidate the syllables into hidden states. To determine the optimal number of hidden states, the log-likelihood using 2-12 states was calculated and determined to be 4 states.

### AlphaClass

Generation of Algorithm: AlphaClass is a supervised behavior segmentation method that employs a similar approach to pose estimation methods to instead detect localized instances of behaviors within an image. Contrary to other existing methods for behavior segmentation that operate on keypoint timeseries (SimBA (110), Keypoint-MoSeq (45)), AlphaClass predicts the likelihood of the presence of behaviors directly from images, similar to DeepEthogram (111). Unlike DeepEthogram, however, AlphaClass operates on single images and predicts the location of where the behavior is occurring by regressing a heatmap, as employed by pose estimation methods for keypoint localization.

Users first begin by defining areas of behavior by placing a point (circle) at the location of a particular behavior, such as placing a point on a grooming mouse, or placing a point at the joint between a mouse’s tail and another mouse’s nose to label chasing behaviors. This is done using makesense.ai. Multiple behaviors can be labeled at a time, such as labeling two mice in a scene that are fighting, while simultaneously labeling a third mouse that is rearing. AlphaClass receives this training data and performs feature detection and keypoint estimation to estimate the likely location of those same user-defined behaviors in untrained frames of video. First, AlphaClass receives an image as an input, such as a single frame from a video of behaving mice. Next, AlphaClass extracts features from these single frames using convolutional neural networks (CNN). The network uses a Resnet50 backbone followed by three upsampling layers, which creates a sigmoid output. This sigmoid output is an input for non-maximum suppression, which is used to identify local peaks and filter our ambiguous detections. The existence of the peaks is interpreted as a positive prediction that the behavior is occurring within the frame whose confidence is the value of the identified peak.

This approach enhances the interpretability of behavior segmentation predictions by providing an estimate of the location where a behavior is occurring, in addition to the confidence. As this bypasses the need for pose estimation, AlphaClass provides a more direct method to detect behaviors that are difficult to classify from pose tracking, such as close social interactions where poses are often noisy due to occlusion. Since AlphaClass only operates on single images, it is limited to classes of behaviors where static image features (e.g., body shape, relative locations) are sufficient to discern whether the behavior is occurring.

Data Analysis: AlphaClass was adapted to quantify the number of “Attending” frames during the Social Exclusion, One Mouse, and Tone Only paradigms, as well as the number of “licking” frames during the formalin assay. Sample frames of the desired behaviors were curated from multiple videos and these frames were labeled using an object detection program, makesense.ai. These frames were then used as training data for AlphaClass and all videos were run through the completed model. A final h5 file was produced for each video and analyzed using MATLAB scripts.

The total number of “Attending” frames was normalized to the baseline number of “Attending” frames present for 5 seconds preceding the onset of the trial. Each 5s interval following trial onset was separately normalized to baseline and total “Attending” frames were calculated by summing the total “Attending” frames over the three time bins from 0-15s.

To characterize the identity of each of the 60 trials and classify them as “Attending” trials or “Not Attending” trials, we applied a global and local threshold analysis using the median value. For the global analysis, the number of “Attending” frames across 60 trials was consolidated across all animals and all paradigms, and the median value of these responses was set as the global threshold. To calculate the local threshold, the number of “Attending” frames across 60 trials was consolidated across each animal for all three paradigms and the median value of these responses was set as the local threshold. Trials that contained more “Attending” frames than this global or local threshold were classified as “Attending” trials, and the rest were classified as “Not Attending” trials. For neural analysis, the local threshold was used per mouse, and the top 15 trials with the most “Attending” frames and bottom 15 trials with the least “Attending” frames were selected to align neural activity to.

### Fiber Photometry

Data acquisition. For fiber photometry experiments, the Neurophotometrics system was used (Neurophotometrics LLC), where a 415 nm LED was used for the reference channel. Frames were captured at 40Hz and each LED was modulated at 20 Hz in an alternating fashion, resulting in a 20Hz sample rate in the reference and signal channels. LED and camera timing as well as recording of timestamps from behavioral equipment was achieved using a data acquisition board (National Instruments NI BNC-2110). Prior to the start of each session, the entire system was shielded from outside light using blackout cloth. Patch cords were obtained from Thor Labs and were photobleached for 24 hours prior to the start of recordings. LED power was calibrated to emit 470 nm light at 50µW for GCaMP activation and 405 nm light at 50µW (measured at the end of the patch cable) through each ferrule, which then interfaced with a ferrule implanted in the mouse (carrying fibers of efficiencies between 80-95%). LEDs were turned on and data was collected continuously for the entire session. Each trial and cue onset were marked through a TTL pulse sent to the Neurophotometrics system.

Data analysis. Both the calcium signal (responses to 470 nm excitation) and reference signals (responses to 405 or 415 nm excitation) were filtered to reject high frequency noise using a forward only median filter with a span of 200ms. Data from the reference channel was then regressed from data in the 470 nm channel. Regression coefficients were computed using data averaged across trials in a session in a time window of −1s to 0s from the start of the CS to minimize any possible regression artifacts introduced by calcium transients recorded in the 470 nm channel evoked by the sensory stimuli or the animal’s response. Residuals from the regression were z-transformed using data from a baseline window of −1s to 0s relative to the start of the CS.

### Cellular Resolution Calcium imaging

Data acquisition: for both Calcium imaging data acquisition and calcium signal extraction nVoke Inscopix systems were used to collect calcium imaging data. During behavior, a TTL signal was used to trigger the miniscope recording to begin. The miniscope was connected to an active commutator (Inscopix). Image processing was accomplished using IDEAS software (Inscopix).

Data analysis: Raw videos were pre-processed by applying 4x spatial downsampling to reduce file size and processing time. A temporal downsampling was applied for a final frame rate of 10Hz. Images were cropped to remove post-registration borders and sections in which cells were not observed. Motion was corrected for by using the first frame as a reference frame. Videos were then exported as TIFF stacks for analysis and converted to an 8-bit TIF file in Fiji ImageJ.

TIFF stacks were then loaded into MATLAB, and additional non-rigid motion correlation was performed using the NoRMCorre package (112). We then used the constrained non-negative matrix factorization algorithm optimized for micro-endoscopic imaging (CNMF-E) (113) to extract fluorescence traces from neurons. Since cellular calcium fluctuations can exhibit negative transients associated with decreases in firing, we did not apply non-negative constraints on temporal components. All neurons were visually confirmed, and neurons exhibiting abnormalities in morphology and calcium traces were excluded. Neuron curation was performed by experimenters blinded to the experimental condition.

To calculate the neuronal response to social and physical pain, the GCaMP7f fluorescence signal for each neuron was z-score normalized to a 1s baseline period immediately preceding the onset of the trial. Trial onset was recorded using the Inscopix system with a single TTL pulse that marked each trial. Z-scores were calculated as (F(t)-Fm)/SD where F(t) is the ΔF/F_0_ at time t and Fm is the mean of ΔF/F_0_ in a baseline period. The z-scored normalized trace was then averaged across a matched number of social trials (“Attending”/”Not Attending”), discrimination trials (reward/shock), and pain trials (0.16gVF, 2.0gVF, pinprick, hot water) for each condition for each neuron. The population mean response was then calculated by averaging the mean z-scored normalized trace of all neurons. The proportion of responsive neurons was calculated using the Wilcoxon signed-rank test.

To co-register neurons between social and physical sessions, we used the cell registration algorithm through an open sourced MATLAB based GUI, CellReg (114). Briefly, spatial footprints of neurons, from CNMF-E output, for all sessions were aligned to a reference session through rotational and translational shifts. Cell pairs were identified by employing a Bayesian probability method that considers the centroid distance between cells and their spatial correlation.

### Decoding

To test if trial types (reward/shock), (“Attending”/”Not Attending”), (dominant, intermediate, subordinate), (0.16gVF, 2.0gVF, pinprick, hot water), could be decoded from single trial aIC population activity, a generalized linear model (GLM) or a support vector machine (SVM) was used. All animals were pooled together to perform global principal component analysis (PCA) over all animals. PCA was only performed on averages of neural activity that would be used for the training set, while omitting the test set. To obtain single trial aIC population activity we used the coefficients obtained for each neuron in the global PCA and created a single trial neural trajectory using the calcium activity for that trial. We used the number of PCs required to explain 90% of the variance, and trained the GLM using these PCs as features, and the trial type as labels. In cases where there were more than two potential categories, as in rank or pain decoding, we used a one-vs-all strategy, where we convert the multi-class problem into four binary primaries (i.e., dominant vs others. intermediate vs others, light touch vs others). We did a 5-fold cross validation for all decoding analyses besides the rank decoding. For these datasets, the data were split into five parts, for 5-fold cross validation and in each iteration the training consisted of a different 80% subset of the data and the testing was done with the remaining 20% of the data. For the rank decoding, we used 10-fold cross validation because more trials were available. For conditions where there were an unequal number of trials, such as reward and shock, where the number of reward trials exceeded shock trials, we randomly subsampled the reward trials to match the number of shock trials for each iteration. We generated the binary classification output in a receiver operating characteristic (ROC) curve, as a function of time. To quantify results, we took the average of the ROC curve over a relevant window, depending on the analysis. For shuffled controls, we repeated the same process with the training data labels shuffled to see the performance of a chance model.

To test if trial type (“Attending” vs “Not Attending”) could be decoded from single trial aIC bulk fluorescence reflecting neuromodulator dynamics for all three of the biosensors used, a random forest (RF) classifier was used. All animals were pooled together and only animals that had successfully undergone all three conditions were included. We did 5-fold cross validation for all decoding, and the data was split into five parts and in each iteration, the training contained 80% of the data and the testing was done on the remaining 20%. All timepoints were pooled together over a relevant window and the ROC obtained using all these data was obtained. For shuffled controls, we repeated the same process using shuffled training data labels.

### Trajectories

To visualize the neural population dynamics in a lower dimensional space, PCA was used to perform dimensionality reduction. A single global PCA was done on a matrix containing all the data for all groups such that neural trajectories could be compared across groups (SE/OM/TO). This matrix had neurons in rows, and in the columns had mean firing rates during −5 to 15s post task-relevant event. All neurons were co-registered across groups so the same neurons would be represented once and horizontally concatenated. The neural trajectories for each task-relevant event were created per group by multiplying the coefficients obtained in the PCA by the mean firing rates across trials.

For each trajectory, the geodesic length was calculated as the sum of Euclidean distances between adjacent 100 timepoints. Distance between trajectories was calculated as the Euclidean distance between the two trajectories bin-by-bin. To allow for statistical comparisons, the neural trajectory metrics were calculated using the leave one out (LOO) method, leaving out all the neurons from a single animal per group, thus the number of iterations is the number of mice in that group. The n reported in trajectory quantifications corresponds to the number of mice utilized for the LOO. Importantly, in every iteration the same PCA coefficients per cell were used for the neural trajectory, since the PCA was done prior to this step, but the neurons included varied. Distance between trajectories was calculated as the Euclidean distance between the two trajectories bin-by-bin. LOO was completed in the same process as for trajectory length calculations. For visualization purposes we plotted the first two or three PC subspaces, but for quantification of trajectory lengths and distance between trajectories the number of PCs that captured 90% of the variance was used. For just the PC space visualizations, we smoothed the trajectories using MATLAB’s smoothdata function.

### Agglomerative hierarchical clustering

Prior to clustering, the data were preprocessed as follows: The peristimulus time histogram (PSTH) was computed using 5s baseline, 15s postcue. Z-scores were calculated using the mean and standard deviation during the baseline period (−1 second to cue onset) individually for each neuron. Data from each experiment protocol (“Attending”, “Not Attending”, and pinprick) were concatenated to calculate universal clusters, allowing for comparisons between each of the social conditions (SE/OM/TO). Using MATLAB, a hierarchical cluster tree was generated using Ward’s method, which uses inner squared distance to determine hierarchy using correlation for the distance metric. A cutoff threshold was used to determine clusters; the value selected was 30% of the maximum value of the linkage distance. Heatmaps plotted for each region are the smoothed z-score input data; clusters for each region are color-coded based on the original cluster tree. All neurons from each cluster were then averaged to create a peri-event time histogram of activity for each cluster during “Attending”, “Not Attending”, and pinprick trials associated with each of our three social sessions. To calculate cluster enrichment, for each social session (SE/TO/OM), the percentage of neurons in each cluster was calculated, and then plotted in comparison with other paradigms.

To determine if a cell was significantly responding to an event, we compared the firing rate in a baseline period vs the event onset (5s window for baseline and event) using a Wilcoxon sign rank test. If firing rate change was significant, excitation or inhibition was determined based on the average z-score during the 15s response window (if it was positive the cell was considered excited, while if it was negative the cell was considered inhibited). The number of responsive neurons was calculated and overlapping neurons responsive to both “Attending”/”Not Attending” and pinprick trials were calculated. Comparisons across different social conditions were done using the chi-squared test. Overlapping excited and inhibited neurons were averaged to plot a per-event time histogram for their responsiveness to pinprick to assess changes in amplitude.

### Histology

Following experiments, mice were deeply anesthetized with sodium pentobarbital (200 mg/kg, intraperitoneal injection). Animals were transcardially perfused with 10 mL of Ringer’s solution followed by 10 mL of cold 4% PFA in 1X PBS. For viral injection, optic fiber, GRIN lens or cannula verification, mice were immediately decapitated after experiments and the whole head was submerged in 4% PFA in 1X PBS for 24 hours at 4°C. The following day the fiber optic implants/GRIN lens/cannulas were removed, and the brains were extracted. Brains were then transferred to 30% sucrose in 1X PBS at 4°C for 12 hours on a shaker. Brains were sectioned coronally at 50µm using a microtome (ThermoScientific) and sections were mounted directly onto glass microscope slides and cover slipped with EMS-Shield Mounting Medium w/ DAPI. Slides were then imaged at 4x magnification using a Keyence BZX710 Fluorescence microscope. Injection sites and implants relative to the mouse atlas were annotated.

### Confocal microscopy

Confocal fluorescence images were acquired on an Olympus FV1000 confocal laser scanning microscope using a 20x/0.75NA objective or a 40x/1.30NA oil immersion objective. Serial Z-stack images were acquired using the FluoView software (Olympus, Center Valley, PA) to confirm viral injections and fiber placements. The tip of the fiber was determined by the 50µm thick gliosis generated by the fiber. The number of cells were quantified with the Imaris software (Bitplane Inc., South Windsor, CT). Regions were located and reported in accordance with the mouse brain atlas.

### Statistics

Statistical analyses were performed using GraphPad Prism 8 (GraphPad Prism, La Jolla, CA) and MATLAB 2023b (Mathworks, Natick, MA). Data with a Gaussian distribution were compared using a paired or unpaired t-test (non-directional) for two experimental groups, and a One-Way or Two-Way ANOVA with repeated measures for three or more experimental groups. To assess responsive neurons, we used a one-sided Wilcoxon rank-sum test. Correlation between two variables was assessed using the Pearson’s correlation coefficient. Kolmogorov-Smirnov statistics were used to detect differences in distribution between two groups. Threshold for significance was placed at p < 0.05. All data are shown as mean ± standard error of the mean (s.e.m).

### Sample size

Sample sizes were based on similar studies in the literature. Sample size is reported in the legends and methods.

### Data exclusions

For calcium experiments, animals were excluded based on histological verification criteria. Histological verification was done by an experimenter blind to the experimental manipulation. Included experimental animals successfully completed 60 trials of SE, OM, and TO, as well as non-nociceptive and nociceptive stimulus application afterwards. Out of eight animals, one animal was excluded based on the histological criteria, and another based on incomplete experimental data collection in Figure 2 and 3. For the Pavlovian discrimination task, one mouse was excluded for learning how to effectively escape from the shock. In photometry experiments, mice were excluded based on histological verification criteria. One animal was excluded for OXT sensor analysis based on lack of expression. One animal was excluded out of OXT sensor OM group analysis based on Grubbs’ test for outliers. For eCB sensor experiments, three animals were excluded for light leak, and seven for lack of expression.

### Reproducibility

Behavioral experiments included in Figure 1 were repeated over two cohorts with multiple investigators. Experiments in Figure 4 were repeated two times.

### Randomization

For behavioral experiments, mice in each cage were randomly divided into SE, OM, and TO groups. Four mice were present in each cage, and each cage had at least one mouse undergoing SE, OM, or TO. The total number of mice undergoing each behavioral paradigm was counter-balanced across mice. For imaging experiments, mice in each cage were randomly divided between experimental and control groups, with two experimental and two control mice in a cage of 4.

### Blinding

During behavioral testing investigators were not always blind to the group affiliation (experimental vs control) given familiarity with the subjects. However, for histology, calcium imaging and biosensor experiments, the experimenters were blinded to the group assignment of the animals (experimental vs control). During data processing and analysis experimenters were blinded to the group affiliation until the point that all data was processed such that group comparisons could be made.

## Data availability

All mouse illustrations included in the main figures were created with BioRender.com. Source data needed to recreate the primary statistical results shown in Figures and Supplementary Figures have been provided as a Supplementary Source Data File. Raw and processed calcium recordings are available on DANDI, and where available we will also include simultaneously recorded behavior videos.

## Code availability

AlphaClass used to process data shown in this manuscript is available on the Tyelab GitHub (https://github.com/Tyelab/AlphaClass).

---

**Extended Data Fig. 1.**
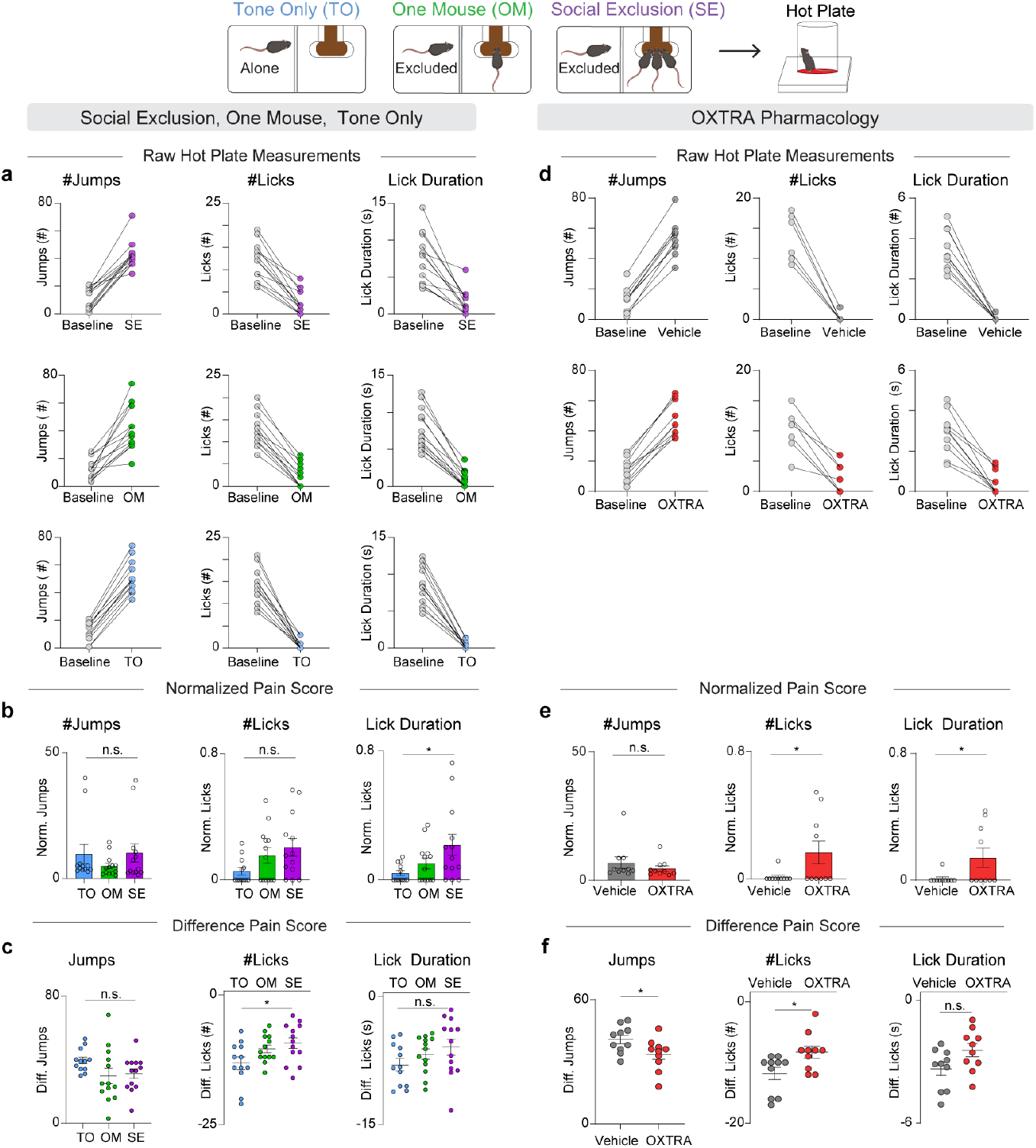
Comparison of different hot plate metrics between experiments. a, Baseline measurements taken before and after SE/OM/TO. These raw values for number of jumps and licks, and lick duration were used to calculate both the normalized pain score and the difference pain score in b and c. b, Normalized readouts for the hot plate relative to baseline conditions before the FOMO Task. Normalized number of jumps during SE/OM/TO (n = 38, One-Way ANOVA F_(2,35)_ = 0.90, p = 0.41). Normalized number of licks during SE/OM/TO (n = 38, One-Way ANOVA, F_(2,35)_ = 2.76, p = 0.08). Normalized lick duration during SE/OM/TO. (n = 38, One-Way ANOVA, F_(2,35)_ = 3.84, *p = 0.031, Tukey’s multiple comparisons test SE vs OM p = 0.15, SE vs TO *p = 0.03, OM vs TO p = 0.35). c, Difference scores for hot plate readouts relative to baseline conditions before the FOMO Task. Difference Score for number of jumps After-Before exposure to the SE/OM/TO (n = 38, One-Way ANOVA F_(2,35)_ = 2.09, p = 0.14). Difference score for number of licks After-Before SE/OM/TO (n = 38, One-Way ANOVA, F_(2,35)_ = 3.85, *p = 0.03, Tukey’s multiple comparisons test, SE vs OM p = 0.72, SE vs TO *p = 0.028, OM vs TO p = 0.14). Difference score for lick duration After-Before SE/OM/TO (n = 38, One-Way ANOVA F_(2, 35)_ = 1.83, p = 0.18). d, Baseline measurements taken before and after SE with intra-aIC OXTRA pharmacology. These raw values for number of jumps and licks and lick duration were used to calculate both the normalized pain score and difference pain score in e and f. e, Responses to hot plate with oxytocin receptor antagonist (OXTRA), L-368,899 hydrochloride at a concentration of 5µg/µl and 200nl per hemisphere into the aIC, relative to vehicle. Normalized number of jumps during OXTRA pharmacology experiments (n = 20 two-tailed unpaired t-test, p = 0.38, t_18_ = 0.91). Normalized number of licks during OXTRA pharmacology experiments (n = 20, two-tailed unpaired t-test, *p = 0.048, t_18_ = 2.12). Normalized lick duration during OXTRA pharmacology experiments (n = 20, two-tailed unpaired t-test, *p = 0.045, t_18_ = 2.16). f, Difference scores for hot plate behavioral measurements. Difference score for number of jumps After-Before exposure to the SE condition paired with intra-aIC OXTRA or vehicle. (n = 20 two-tailed unpaired t-test, *p = 0.04, t_18_ = 2.25) Difference score for number of licks upon intra-aIC OXTRA administration. (n = 20, two-tailed unpaired t-test, *p = 0.02, t_18_ = 2.49). Difference score for lick duration After-Before exposure to the SE condition paired with intra-aIC OXTRA or vehicle. (n = 20, two-tailed unpaired t-test, p = 0.06, t_18_ = 2.04).

**Extended Data Fig. 2.**
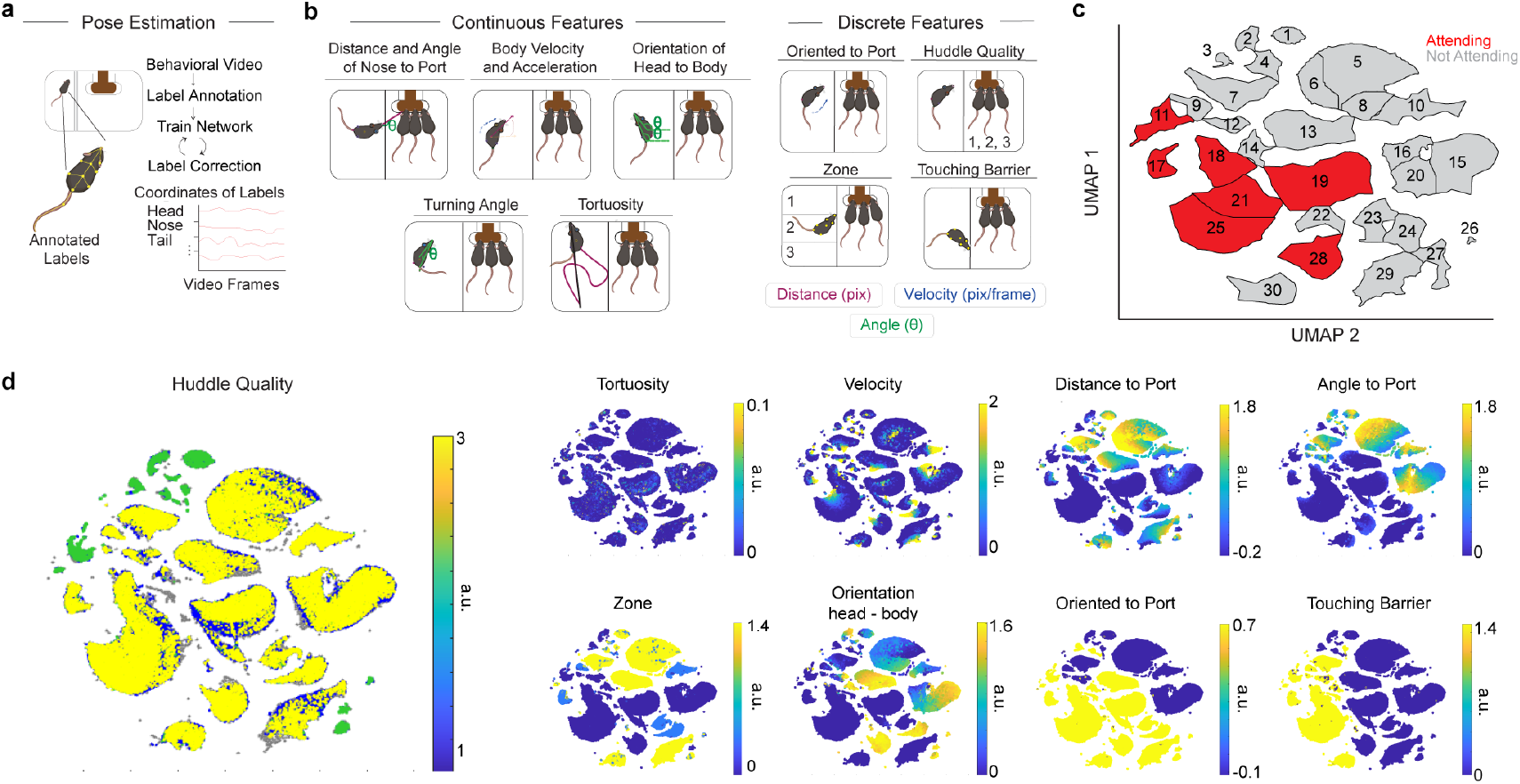
Distinct clusters are discovered through UMAP and DBSCAN. a, Mice poses were estimated using SLEAP. b, The SLEAP model was iteratively trained using discrete features that were extracted as behavioral features. Tortuosity describes the path that the animal takes over 1s. Orientation to port is either classified as 0 or 1, depending on angle to port. Huddle Quality is classified as the number of mice at the port at any time (1, 2, 3). Zone is classified as the zone that the excluded mouse is in (1, 2, 3). Touching Barrier is classified as 0 or 1, depending on the distance to the barrier. Colors indicate different metrics, including distance in pixels, velocity, and angle. c, DBSCAN clusters on the UMAP. Each cluster is labeled, and colors indicate whether that cluster was manually selected to be an “Attending” cluster or an “Not Attending” cluster. d, Features were mapped onto UMAP to identify how clusters were influenced by each behavioral feature.

**Extended Data Fig. 3.**
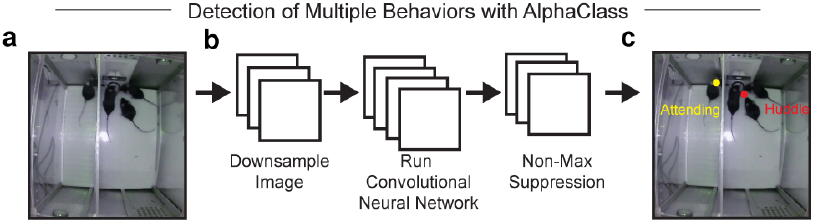
AlphaClass Architecture. a, Model architecture for AlphaClass, modified from (119). AlphaClass is a 2-D behavioral estimate method that primarily uses keypoint detection to estimate behaviors in single images. b, For training, AlphaClass receives images as input and extracts features through a downsampling method with a convolutional neural network (CNN). Then, non-max suppression is used to remove any faulty detections or low-confidence detections. c, The final results are behavioral labels predicted on novel video data. Example of final behaviors used for analysis during Social Exclusion. “Attending” frames were quantified when the excluded mouse was labeled as “Attending”.

**Extended Data Fig. 4.**
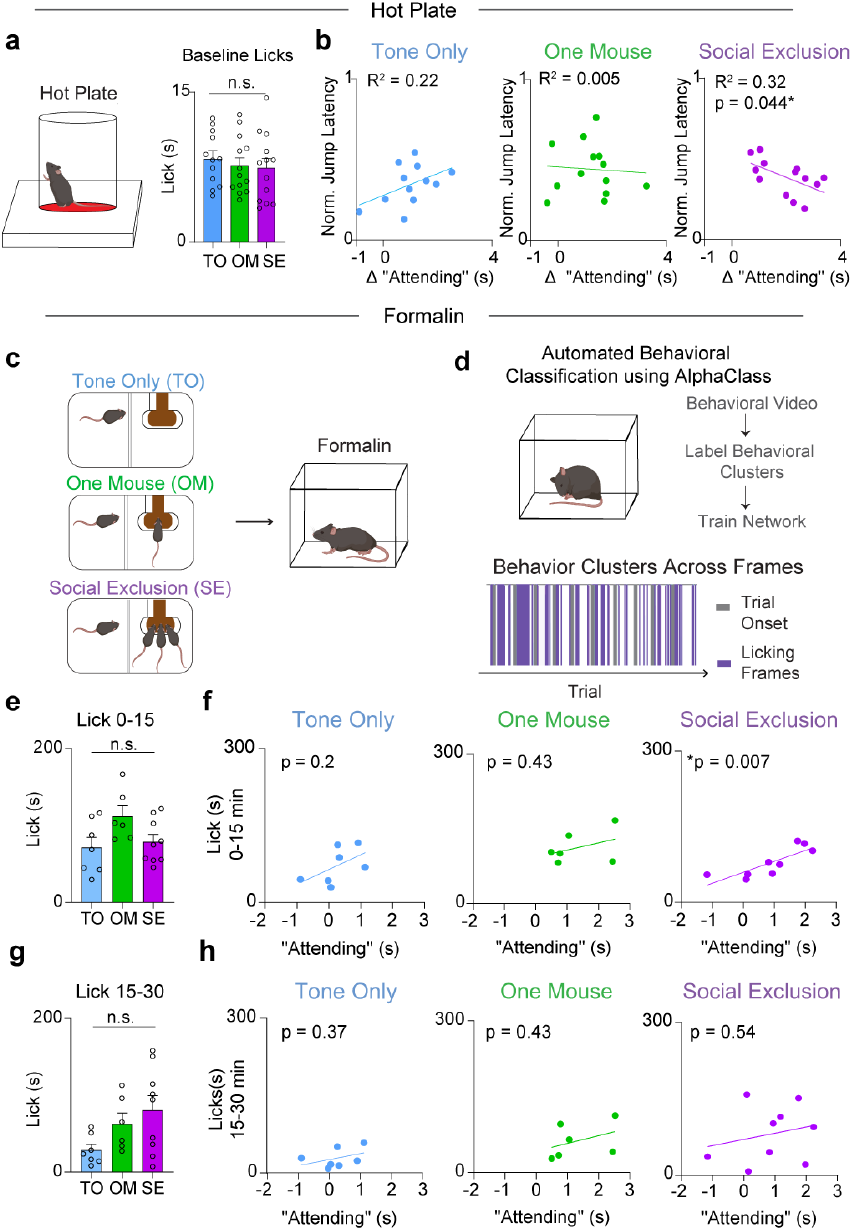
Social Exclusion “Attending” behavior is correlated with physical pain. a, Baseline measurements for each mouse on the hot plate. The paradigm group that each animal is placed in is the future social paradigm that that animal will experience. (n = 38, One-Way ANOVA, F_(2, 35)_ = 0.33, p = 0.72). b, Correlation between “Attending” frames and behavioral responses to Physical Pain. TO (n = 12, Linear Regression R^2^ = 0.22, p = 0.1226), OM (n = 13, Linear Regression R^2^ = 0.005, p = 0.82), SE (n = 13, Linear Regression R^2^ = 0.32, *p = 0.04) c, Mice underwent one of three behavioral conditions (SE/OM/TO) and then were injected with formalin into the right hindpaw. d, AlphaClass was used to quantify lick duration and bouts. e, Lick duration after formalin injection during the first phase. There is no difference between duration of licks during the first phase after formalin injection. (n = 23, One-Way ANOVA, F_(2,19)_ = 0.20. p = 0.82). f, Correlation between licks during the first phase after formalin injection, and “Attending” behavior. Mice that exhibit more “Attending” behavior during Social Exclusion lick more after formalin injection. TO (n = 7, Linear Regression, R^2^ = 0.30, p = 0.20). OM (n = 6, Linear Regression R^2^ = 0.16, p = 0.43). SE (n = 9, Linear Regression, R^2^ = 0.67, **p = 0.007). g, Lick duration after formalin injection during the second phase. There is no difference between groups (n = 23, One-Way ANOVA, F_(2,19)_ = 3.8, p = 0.07). h, Correlation between licks during the second phase after formalin injection and “Attending” behavior. There is no correlation in any of the behavioral conditions. TO (n = 7, Linear Regression R^2^ = 0.16, p = 0.37). OM (n = 6, Linear Regression R^2^ = 0.16, p = 0.43). SE (n = 9, Linear Regression R^2^ = 0.06, p = 0.54).

**Extended Data Fig. 5.**
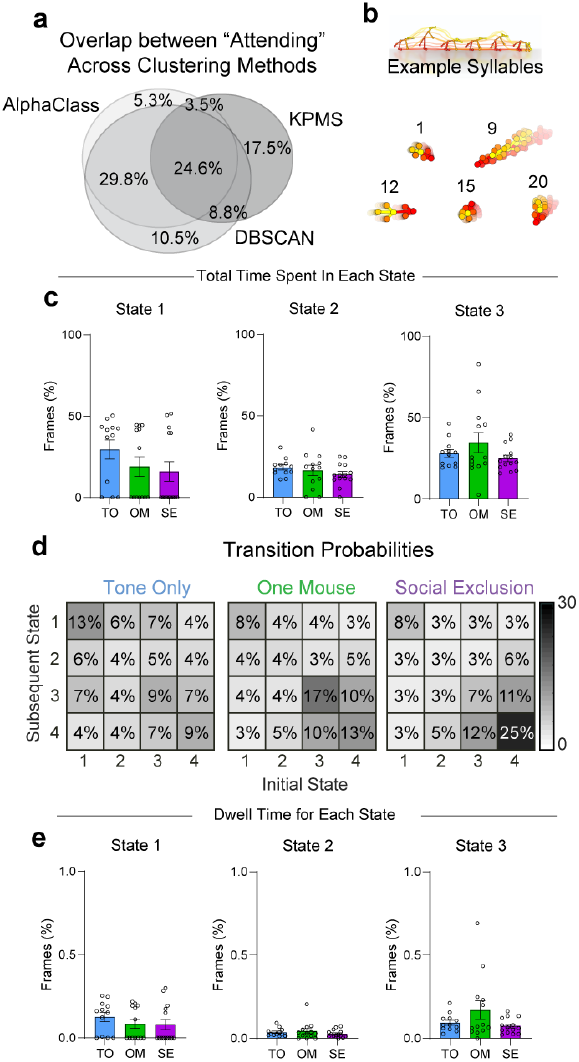
Additional behavioral metrics for other hidden states during the “FOMO” task. a, Venn Diagram displaying percentage of overlapping frames across the three supervised and unsupervised behavioral classification methods used. b, Example syllables generated using KeypointMoseq (KPMS) that are present in the FOMO task. A total of 87 syllables were generated using all frames from the Social Exclusion, One Mouse, and Tone Only paradigms with n = 39. c, Duration spent in each hidden state between social conditions. Besides State 4, there is no difference for all states between conditions. (n = 39, One-Way ANOVA, State 1 F_(2,36)_ = 0.11, p = 0.27, State 2 F_(2,36)_ = 2.18, p = 0.43. State 3 F_(2,36)_ = 3.58, p = 0.23). d, Transition Probabilities between each of the four behavioral states. Transitions are grouped over all animals per condition and normalized across the entire matrix. e, Dwell Time for each hidden state per social condition. Besides State 4, there is no difference between behavioral conditions for each state. (n = 39, One-Way ANOVA, State 1 F_(2,36)_ = 0.02, p = 0.49. State 2 F_(2,36)_ = 1.03, p = 0.53, State 3 F_(2,36)_ = 2.93, p = 0.12). Error bars around the mean indicates s.e.m.

**Extended Data Fig. 6.**
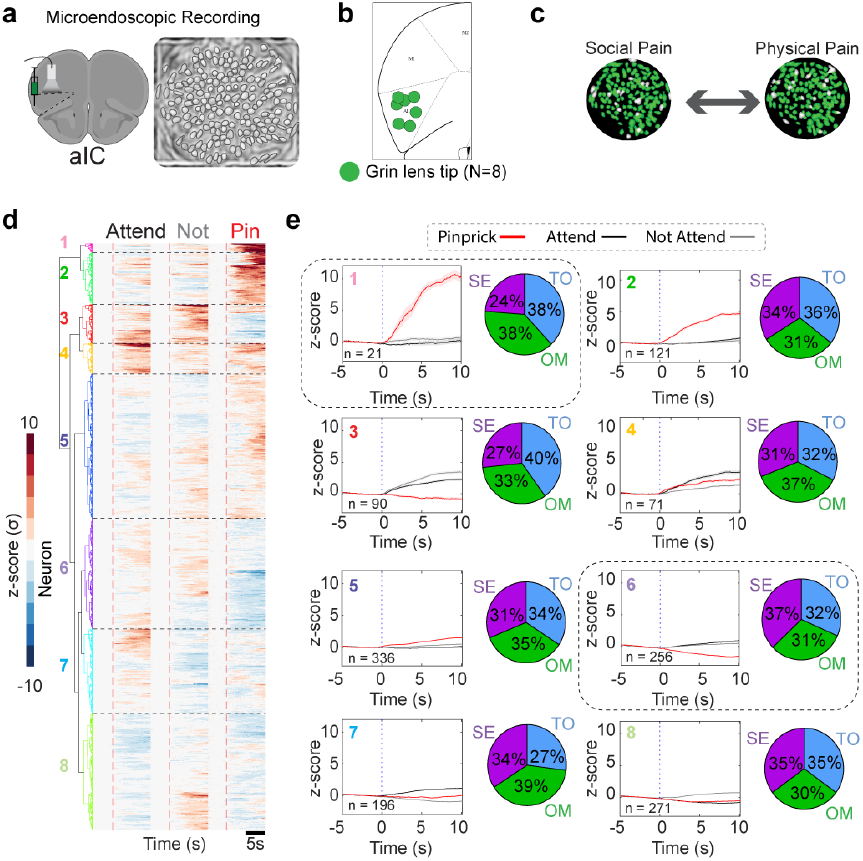
Social Exclusion causes distinct responses to pinprick. a, Schematic of viral strategy for cellular resolution calcium imaging of anterior insular cortex (aIC) cells. Example image of contours extracted using constrained non-negative matrix factorization for microendoscopic data (CNMF-e). Each outline represents one neuron. b, Location of GRIN lens (0.5 mm x 0.67 mm) implant within the aIC for all animals used across both social and discrimination single unit analysis. c, Example output from cell registration to coregister and track the same cells across social and physical pain stimuli. d, Neuron responses are separated into clusters using hierarchical clustering. Heatmap for aIC ΔF/F_0_ responses to “Attending” and “Not Attending” trials during the Social Exclusion, One Mouse, and Tone Only paradigms, as well as during pinprick stimuli applied after each social condition. Social trials are aligned to trial onset. At trial onset, the cue is played, and the switchable glass wall turns transparent. Pain trials are aligned to stimulus application onset. Neurons are co-registered per social condition and physical pain stimuli application. Colors indicate clusters obtained using hierarchical clustering. e, Response amplitude for aIC cells during “Attending”, “Not Attending”, and pinprick trials for each specific cluster. Cluster colors are matched with heatmap cluster (d). Pie charts depict the percentage of neurons that belong to each social condition.

**Extended Data Fig. 7.**
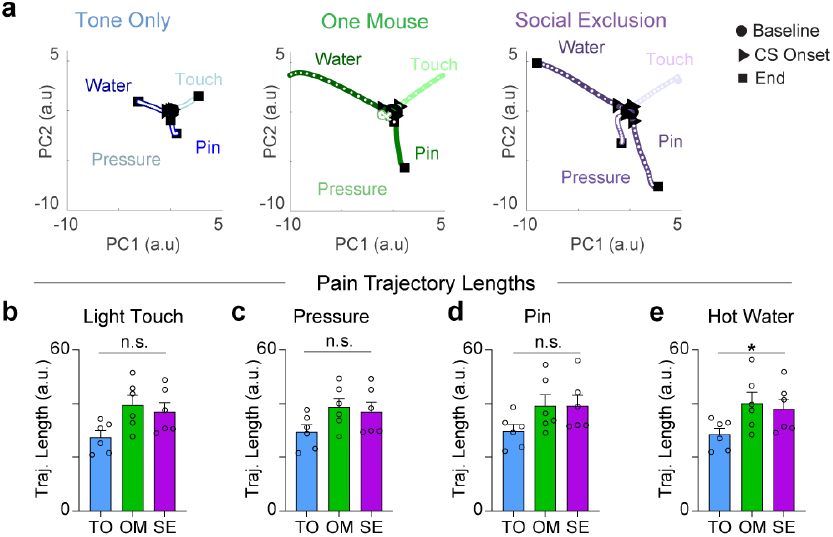
Neural trajectories from aIC neural population responses to physical pain stimuli. a, Sample neural trajectories for one representative animal. Responses to light touch, pressure, pinprick, and hot water are shown and graded by color to match the stimulus intensity. b-e, Quantification of trajectory lengths. b, Touch: Repeated Measures One-Way ANOVA, n = 6, F_(1.6,8.2)_ = 4.6, p = 0.051. c, Pressure: Repeated Measures One-Way ANOVA, n = 6, F_(1.5,7.2)_ = 2.74, p = 0.136. d, Pin: Repeated Measures One-Way ANOVA, n = 6, F_(1.7,8.5)_ = 3.1, p = 0.098. e, Water: Repeated Measures One-Way ANOVA, n = 6, F_(1.9,9.4)_ = 4.3, *p = 0.049.

**Extended Data Fig. 8.**
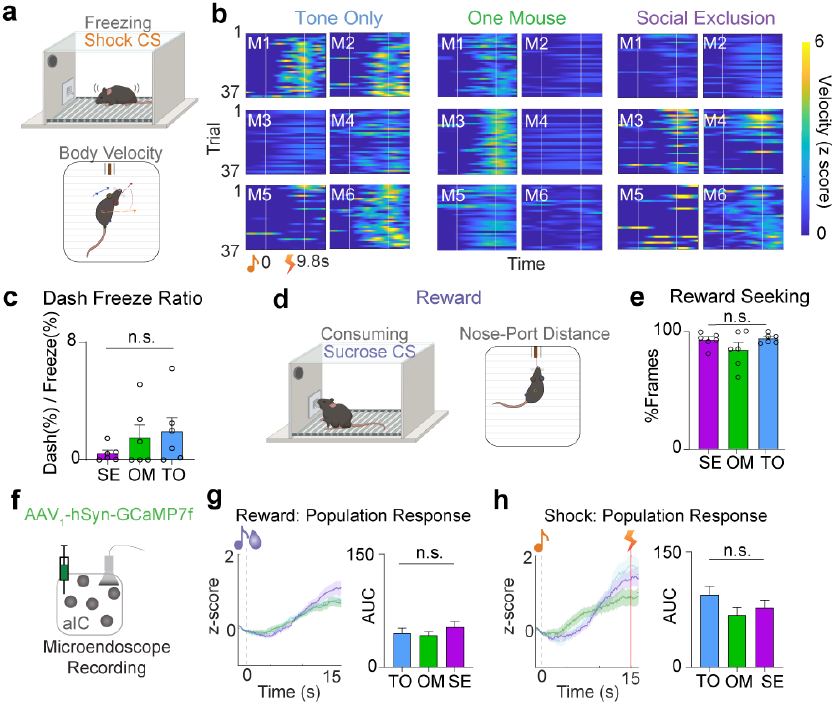
Social Exclusion only changes encoding of conditioned cues, not behavioral responses to reward and shock. a, Quantification of freezing behavior during shock trials using body velocity. Velocity was z-scored for each animal separately across its three sessions. b, Heatmaps showing normalized mouse velocity for 37 shock trials (n = 6). Cue onset and shock onset are marked at 0s and 9.8s respectively. c, Ratio of dash/freezing behavior for each animal across social conditions in the 0-15s post-CS onset (Repeated Measure One-Way ANOVA, F_(1.1,5.4)_ = 0.89. p = 0.39). d, Quantification of reward seeking behavior during reward trials using nose-port distance. A threshold was set for all animals for distance to the port to determine number of reward frames. e, Percentage of frames under the nose-port distance threshold to quantify reward seeking. There is no difference in reward seeking behavior across social conditions (Repeated Measures One-Way ANOVA, F_(1.4,6.8)_ = 1.39, p = 0.30) f, Viral strategy for labeling aIC neurons with a calcium indicator to record activity during the discrimination assay. g,h, Average population neural activity for both reward (g) and shock trials (h). All responsive neurons were determined using rank-sum for each neuron. Final responsive neurons were averaged for a population response to both trial types. g, There is no difference in the average z-score responses to reward trials after the three social conditions. (n = 6, One-Way ANOVA reward: F_(2,278)_ = 0.76, p = 0.47. h, There is no difference in responses to shock. (n = 6, One-Way ANOVA shock: F_(2,228)_ = 1.59, p = 0.21). Error bars and solid shaded regions around the mean indicates s.e.m.

**Extended Data Fig. 9.**
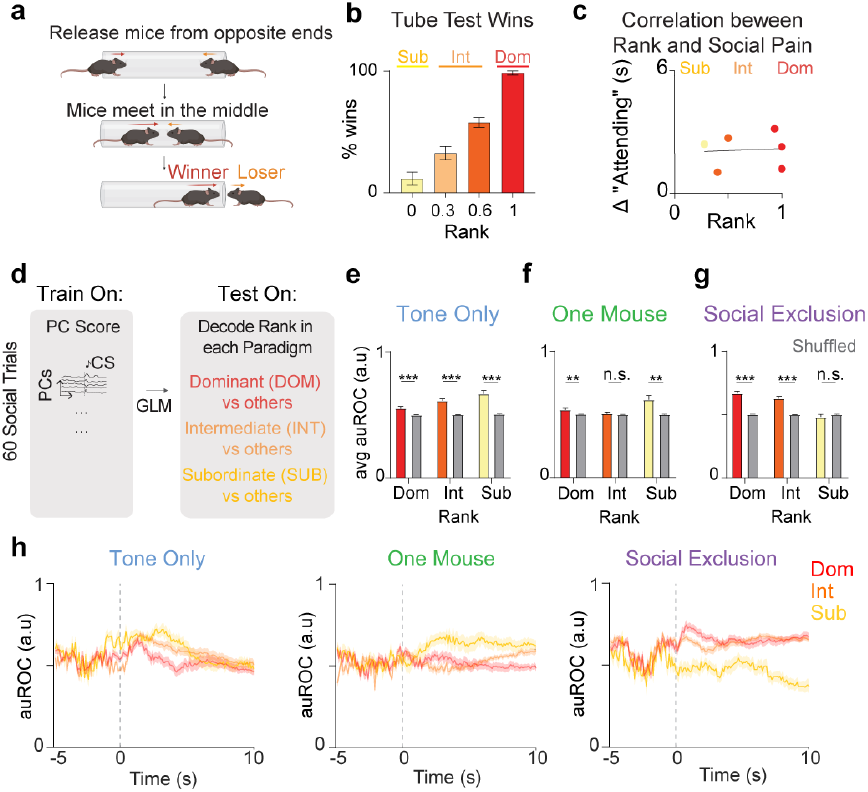
aIC rank encoding dynamically shifts based on FOMO Task Condition. a, Tube test schematic to determine rank. Mice are tested in a round robin fashion. b, Rank is determined by overall percentage of wins during the four days of testing (n = 16, One-Way ANOVA, F_(3,12)_ = 71.62, ***p<0.0001. Tukey’s multiple comparisons test. Rank 1 vs 2, ***p = 0.0001, Rank 1 vs 3 ***p<0.0001, Rank 1 vs 4, ***p < 0.0001. Rank 2 vs 3, **p = 0.0075, Rank 2 vs 4, ***p<0.0001, Rank 3 vs 4 *p = 0.026). c, Correlation between Social Rank and “Attending” frames. There is no relationship between social rank and the amount of “Attending” behaviors exhibited (n = 6, Pearson’s Correlation, R^2^ = 0.005, p = 0.89). d, Decoding schematic using a generalized linear model to test if rank information is represented in neural responses to 60 social trials. The number of PCs required to explain 90% of the variance was used in this analysis, and decoding was done with 10-fold cross validation. e, Area under the ROC (auROC) for each rank in the Tone Only paradigm. All ranks are decodable during Tone Only (n = 6, Two-Way ANOVA, Main effect Rank F_(2,354)_ = 38.3, *** p < 0.0001, Shuffled Control F_(1,354)_ = 358, *** p < 0.0001, Rank x Shuffled Control F_(2,354)_ = 31.58, ***p < 0.0001. Tukey’s multiple comparisons test. Dom vs Dom Shuffled ***p < 0.0001. Int vs Int Shuffled ***p<0.0001. Sub vs Sub Shuffled ***p < 0.0001. Tukey’s multiple comparisons test. Sub vs Int ***p < 0.0001, Sub vs Dom ***p < 0.0001, Int vs Dom ***p<0.0001). f, Area under the ROC (auROC) for each rank in the One Mouse paradigm. Only Dominants and Subordinates are decodable during One Mouse within the aIC (n = 6, Two-Way ANOVA, Main effect Rank F_(2,354)_ = 24.85, ***p < 0.0001, Shuffled Control F_(1,354)_ = 63.8, ***p < 0.0001, Rank x Shuffled Control F_(2,354)_ = 24.1, ***p < 0.0001. Tukey’s multiple comparisons test. Dom vs Dom shuffled **p = 0.009. Int vs Int shuffled p = 0.86. Sub vs Sub shuffled ***p < 0.0001. Tukey’s multiple comparisons test Dom vs Sub ***p < 0.0001. Dom vs Int p = 0.15. Int vs Sub ***p<0.0001. g, Area under the ROC (auROC) for each rank in the Social Exclusion paradigm. Only Dominants and Intermediates are decodable during Social Exclusion within the aIC. (n = 6, Two-Way ANOVA, Main effect Rank F_(2,354)_ = 97.94, ***p < 0.0001, Shuffled Control F_(1,354)_ = 257.5, ***p < 0.0001, Rank x Shuffled Control F_(2,354)_ = 97.3, ***p < 0.0001. Tukey’s multiple comparisons test. Dom vs Dom shuffled ***p < 0.0001. Int vs Int shuffled ***p<0.0001. Sub vs Sub shuffled p = 0.17. Tukey’s multiple comparisons test Dom vs Sub ***p < 0.0001. Dom vs Int **p = 0.004. Int vs Sub ***p<0.0001. h, Decoding performance of rank displayed in 100 ms bins for each social condition. Error bars and solid shaded regions around the mean indicates s.e.m.

**Extended Data Fig. 10.**
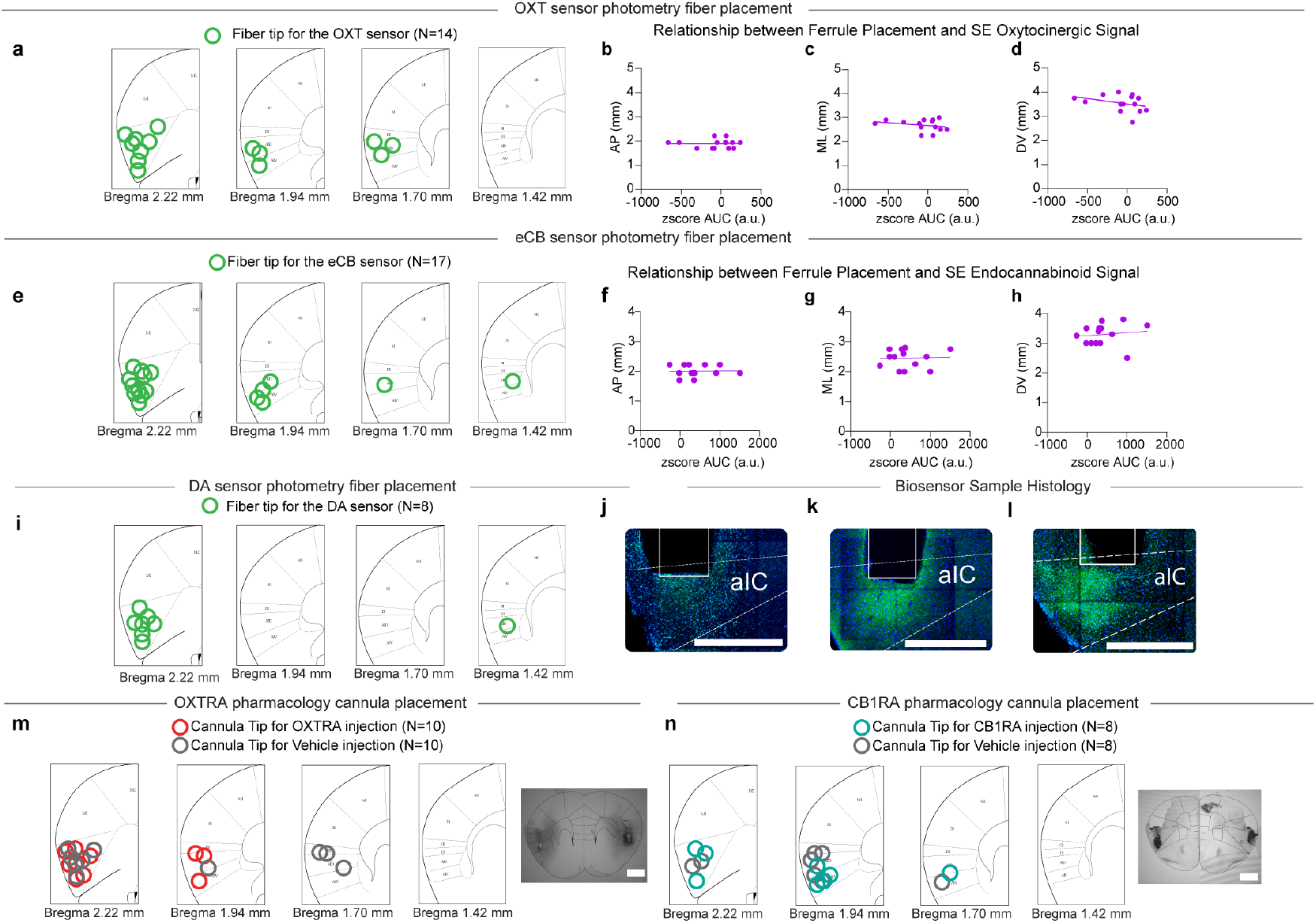
Biosensor and Pharmacology Verification. a, Histological verification of ferrule placement for the oxytocin sensor experimental cohort. Each green dot marks the tip of the ferrule for each animal. One animal was excluded due to lack of expression. b-d, Correlation between the oxytocinergic signal during Social Exclusion, and placement of ferrule. Anterior-posterior (AP), Medial-lateral (ML), Dorsal-ventral (DV). b, Correlation with anterior-posterior coordinates R^2^ = 0.0002, p = 0.96. c, Correlation with medial-lateral coordinates R^2^ = 0.07, p = 0.37. d, Correlation with dorsal ventral coordinates R^2^ = 0.12. p = 0.23. e, Histological verification of ferrule placement for the endocannabinoid sensor experimental cohort. Each green dot marks the tip for each animal. Three animals were excluded for light leak. Seven animals were excluded for lack of expression. f-h, Correlation between the endocannabinoid signal during Social Exclusion and placement of ferrule. f, Correlation with anterior-posterior coordinates R^2^ = 0.0003, p = 0.95, g, Correlation with medial-lateral coordinates R^2^ = 0.001, p = 0.91. (h) Correlation with dorsal-ventral coordinates R^2^ = 0.017, p = 0.66. i, Histological verification of ferrule placement for the dopamine sensor experimental cohort. No animals were excluded. j, Example histology image of GRAB_OXT1.0_ sensor expression. Scale Bar 500 µm. k, Example histology image of GRAB_eCB2.0_ sensor expression. Scale Bar 500µm. l, Example histology image of GRAB_DA3h_ sensor expression. Scale Bar 500µm. m, Histological verification of cannula placement for the OXTRA pharmacology experiments. Three animals were excluded based on cannula placement. Colors indicate whether the animal received an OXTR antagonist or Vehicle Injection. Scale Bar 1000µm. n, Histological verification of cannula placement for the CB1R agonist pharmacology experiments. No animals were excluded based on cannula placement. Scale Bar 1000µm.

